# mRNA structure regulates protein expression through changes in functional half-life

**DOI:** 10.1101/549022

**Authors:** David M. Mauger, B. Joseph Cabral, Vladimir Presnyak, Stephen V. Su, David W. Reid, Brooke Goodman, Kristian Link, Nikhil Khatwani, John Reynders, Melissa J. Moore, Iain J. McFadyen

**Author notes:** Co-corresponding Authors: Melissa J. Moore and Iain J. McFadyen. Co-lead Contact: Melissa J. Moore and Iain J. McFadyen.

## Abstract

Messenger RNAs (mRNAs) encode information in both their primary sequence and their higher order structure. The independent contributions of factors like codon usage and secondary structure to regulating protein expression are difficult to establish as they are often highly correlated in endogenous sequences. Here, we used two approaches, global inclusion of modified nucleotides and rational sequence design of exogenously delivered constructs to understand the role of mRNA secondary structure independent from codon usage. Unexpectedly, highly-expressed mRNAs contained a highly-structured coding sequence (CDS). Modified nucleotides that stabilize mRNA secondary structure enabled high expression across a wide-variety of primary sequences. Using a set of eGFP mRNAs that independently altered codon usage and CDS structure, we find that the structure of the CDS regulates protein expression through changes in functional mRNA half-life (i.e. mRNA being actively translated). This work highlights an underappreciated role of mRNA secondary structure in the regulation of mRNA stability. [150 words]

**Highlights:**

- Protein expression from modified mRNAs tends to follow the pattern m^1^ Ψ > U >mo^5^U
- Protein expression correlates with mRNA thermodynamic stability: Ψ≈ m^1^Ψ > U > mo^5^U
- A highly structured CDS correlates with high expression
- Increased structured mRNAs extend functional half-life

## Introduction

Messenger RNAs (mRNAs) direct cytoplasmic protein expression. How much protein is produced per mRNA molecule is a function of how well the translational machinery initiates and elongates on the coding sequence (CDS), combined with the mRNA’s functional half-life. Both translational efficiency and functional half-life are driven by features encoded in the primary mRNA sequence. Synonymous codon choice directly impacts translation, with highly expressed genes tending to include more “optimal” codons (Gustafsson et al., 2004; Hinnebusch et al., 2016; Horstick et al., 2015; Pop et al., 2014). Conversely, “non-optimal” codons can increase ribosomal pausing and decrease mRNA half-life (Presnyak et al., 2015; Weinberg et al., 2016). Other mRNA sequence features reported to correlate with protein output are dinucleotide frequency in the CDS (Tulloch et al., 2014), and the effect of codon order on locally-accessible charged tRNA pools (Tuller et al., 2010). Because these effects are interdependent on mRNA sequence, teasing apart their individual contributions to protein output are difficult and often controversial (Futcher et al., 2015; Simmonds et al., 2015).

In addition to dictating encoded protein identity, its primary sequence also determines an mRNA’s propensity to form secondary and tertiary structure (Mortimer et al., 2014). Transcriptome-wide RNA structure characterization is beginning to reveal global relationships between the structure content in different mRNA regions and protein expression (Ding et al., 2014; Kertesz et al., 2010; Ramani et al.; Wan et al., 2014). For example, multiple studies across all kingdoms of life, from bacteria to humans, have shown that secondary structure in the 5’ untranslated region (5′UTR) generally reduces translation initiation efficiency and therefore overall protein output (Ding et al., 2014; Gu et al., 2010; Shah et al., 2013; Tuller and Zur, 2015; Wan et al., 2014). The extent to which CDS and 3’ untranslated region (3′UTR) secondary structure impacts protein output, however, is much less understood.

One way to alter RNA secondary structure is to change the primary sequence. In the CDS, however, primary sequence changes necessarily alter codon usage, confounding any effects that might be attributable to changes in mRNA structure alone. An alternate means to affect secondary structure without changing codons is to incorporate modified nucleotides that maintain the same Watson-Crick base pairing relationships (e.g., pseudouridine (Ψ) for U) and have small effects on local secondary structure. Such modified nucleotides can either stabilize (Newby and Greenbaum, 2001) or destabilize (Kierzek and Kierzek, 2003) base pairs and hence overall mRNA structure. By destabilizing local secondary structure in endogenous mRNAs, and thereby increasing the regulatory protein accessibility to specific sequence elements, N1-methyl and N6-methyl adenosine are emerging as key regulators of protein expression (Dominissini et al., 2016; Spitale et al., 2015). Other modified nucleotides known to exist within mRNA and affect secondary structure formation include Ψ, 5-methyl-cytidine, and inosine (Harcourt et al., 2017).

Recent progress in synthesizing and delivering exogenous mRNAs (Kariko et al., 2008; Sabnis et al., 2018) opens the possibility of more broadly exploring the relationship between mRNA structure and protein output by comparing expression of mRNAs having identical sequences but different modified nucleotide content. Here, we combined computational sequence design with global modified nucleotide substitution as tools to investigate the separate impacts of mRNA primary sequence and structural stability on protein output. We did so across multiple synonymous mRNA sequence variants encoding three different proteins. We find that differences in the innate thermodynamic base pair stability of two modified uridine nucleotides, N1-methyl-pseudouridine and 5-methoxy-uridine, induce global changes in mRNA secondary structure. These structural changes in turn drive changes in protein expression. As expected, our data confirm that reduced secondary structure within a 5’ leader region (the 5’ UTR and first ~10 codons of the CDS) correlates with high protein expression. Surprisingly, we also find that high protein expression correlates with increased secondary structure in the remainder of the mRNA (the rest of the CDS and the 3’ UTR). We validated this finding by designing an eGFP mRNA panel wherein the effects of codon usage and secondary structure could be examined separately. Our data reveal a relationship wherein codon optimality and greater CDS secondary structure synergize to increase mRNA functional half-life. Thus, by decoupling structural stability from primary sequence changes, exogenously provided mRNAs containing modified nucleotides provide a valuable tool to specifically investigate the contributions of mRNA structure to protein expression.

## Results

### RNA sequence and nucleotide modifications combine to determine protein expression

For this study, we created diverse synonymous CDS sets encoding enhanced Green Fluorescent Protein (eGFP, four variants), human erythropoietin (hEpo, nine variants) and firefly luciferase (Luc, 39 variants) transcribed *in vitro* with ATP, CTP, GTP and either UTP, pseudouridine triphosphate (ΨTP), N^1^-methyl-pseudouridine triphosphate (m^1^ΨTP), or 5-methyoxy-uridine triphosphate(mo^5^UTP) (Figure 1A). For comparison with a previous study documenting the effects of modified nucleotides on RNA immunogenicity (Kariko et al., 2008), we also made eGFP mRNA wherein both U and C were substituted with Ψ and 5-methyl-cytidine (m^5^C), respectively. We designed the sequence sets with bias towards “optimal” codons (for hEPO and eGFP mRNAs) or designed to sample a larger sequence space, including “non-optimal” codons (for Luc mRNA). All mRNAs carried a 7-methylguanylate cap (m^7^G-5’ppp5’-Gm), identical 5ʹ and 3ʹ UTRs, and a 100-nucleotide poly(A) tail.

**Figure 1.**
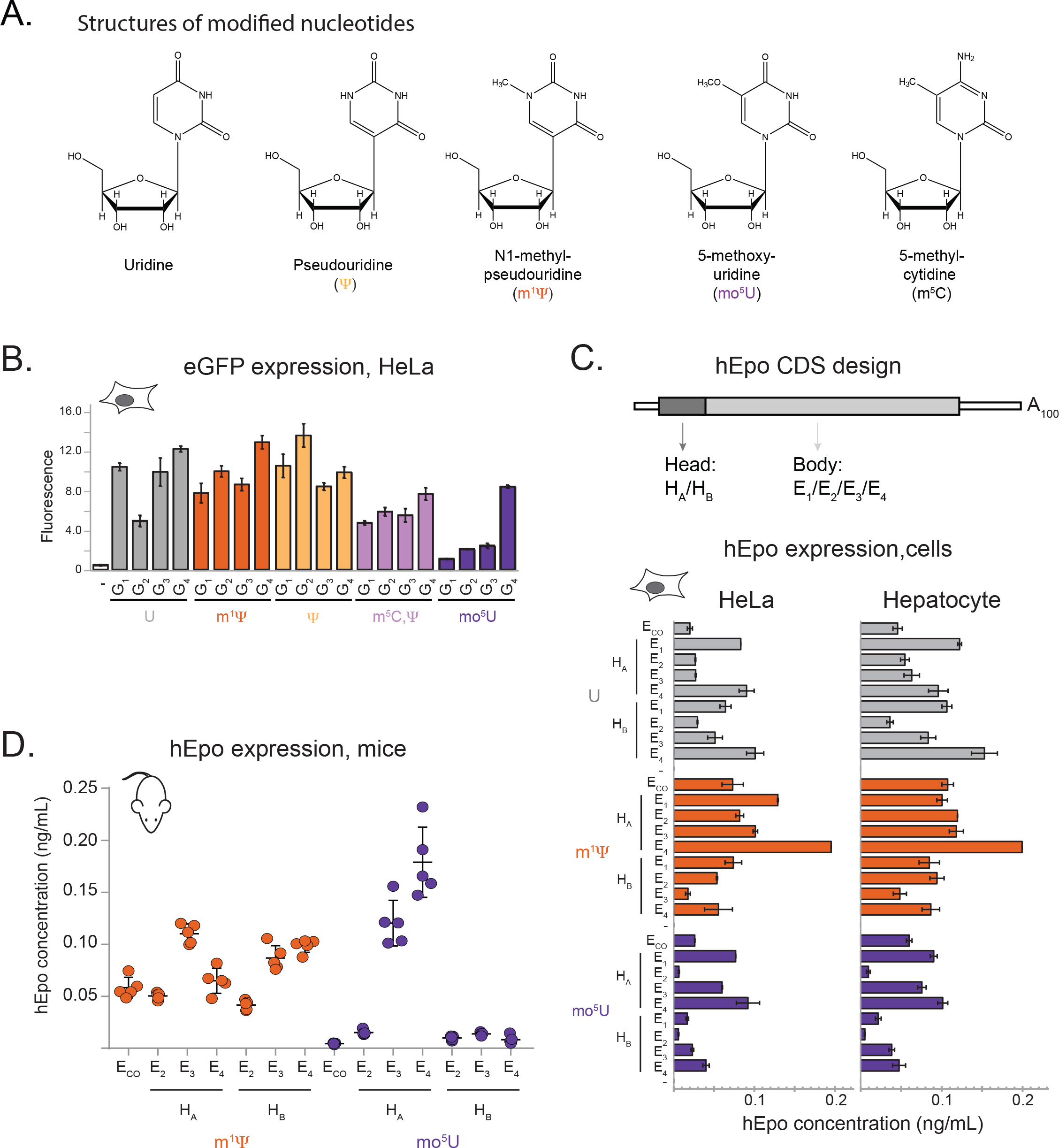
Inclusion of modified nucleotides in mRNA alters eGFP and hEPO expression. (A) Chemical structure of uridine and four modified nucleosides: pseudouridine (Ψ), N^1^-methyl-pseudouridine (m^1^Ψ), 5-methoxy-uridine (mo^5^U), and 5-methyl-cytidine (m^5^C). (B) Fluorescence intensity (normalized intensity units, y-axis) of HeLa cells following transfection with lipofectamine alone (-) or four different eGFP sequence variants (G_1_-G_4_, x-axis) containing uridine (grey), m^1^Ψ (orange), Ψ (yellow), m^5^C/Ψ (lavender), or mo^5^U (dark purple). (C) Top panel: schematic of the human erythropoietin (hEpo) mRNA sequence variants. The coding sequence (wide grey boxes) is flanked by 5’and 3’untranslated regions (UTRs, narrow white boxes) and a 3ʹ 100-nucleotide poly-A tail. Eight hEpo sequences combined one of two “head” regions (dark grey box, H_A_ or H_B_) encoding the first 30 amino acids (90 nucleotides) and one of four “body” regions (light grey box, E_1_ through E_4_) encoding the remainder of the hEpo CDS. Bottom panel: levels of secreted hEpo protein measured by ELISA (ng/mL, y-axis) following transfection of cells with 8 sequence variants (described in B above, x-axis) plus one “codon optimized” variant (E_CO_) (Welch et al., 2009) containing uridine (grey bars), m^1^Ψ (orange), or mo^5^U (dark purple). (D) Serum concentrations of hEpo protein measured by ELISA (ng/mL, y-axis) in BALB-c mice (five per group) following IV injection of LNP-formulated mRNA of 6 sequence variants (described in B above, x-axis) plus one “codon optimized” variant (E_CO_) (Welch et al., 2009) containing m^1^Ψ (orange) or mo^5^U (dark purple). Individual animals (dots) with mean and standard error (black lines).

First, we analyzed the impact of primary CDS sequence on protein expression of mRNAs containing no modified nucleotides (eGFP/hEPO, Figure 1; Luc, Figure 2). Cellular protein expression ranged >2.5-fold for eGFP (Figure 1B, grey) and >4-fold for hEpo (Figure 1C, grey), despite all sequences containing only frequently used codons. Expression of 39 unmodified Luc variants containing codons with a greater optimality range varied >10-fold (Figure 2A). Highly expressed mRNAs tended to have increased GC content, consistent with previous reports (Plotkin and Kudla, 2011), but not all high GC sequences were high expressers (Figure S1 A & B, Figure S2A, grey). Unmodified Luc variant expression moderately correlated with both GC-content and Codon Adaptation Index (CAI) (Pearson correlations *r* =0.63 and 0.64, respectively, Figure S2A, grey). Each Luc variant globally used the same single codon for all instances of a given amino acid. This allowed us to look at the impact of individual codons on protein expression. Only 4 of 87 pairwise synonymous codon comparisons exhibited statistically significant differences (*p* < 0.05, Figure S3, grey). For example, inclusion of Phe^UUU^ was associated with a slight increase in expression over Phe^UUC^ (Figure 2B). Surprisingly, even global inclusion of extremely non-optimal codons such as Ser^UCG^, Leu^CUA^, Ala^GCG^, and Pro^CCG^ had no statistically significant impact on Luc expression in unmodified RNA (Figure 2B, Figure S3). Thus, codon usage, as measured by metrics such as CAI, cannot adequately explain these data.

**Figure 2.**
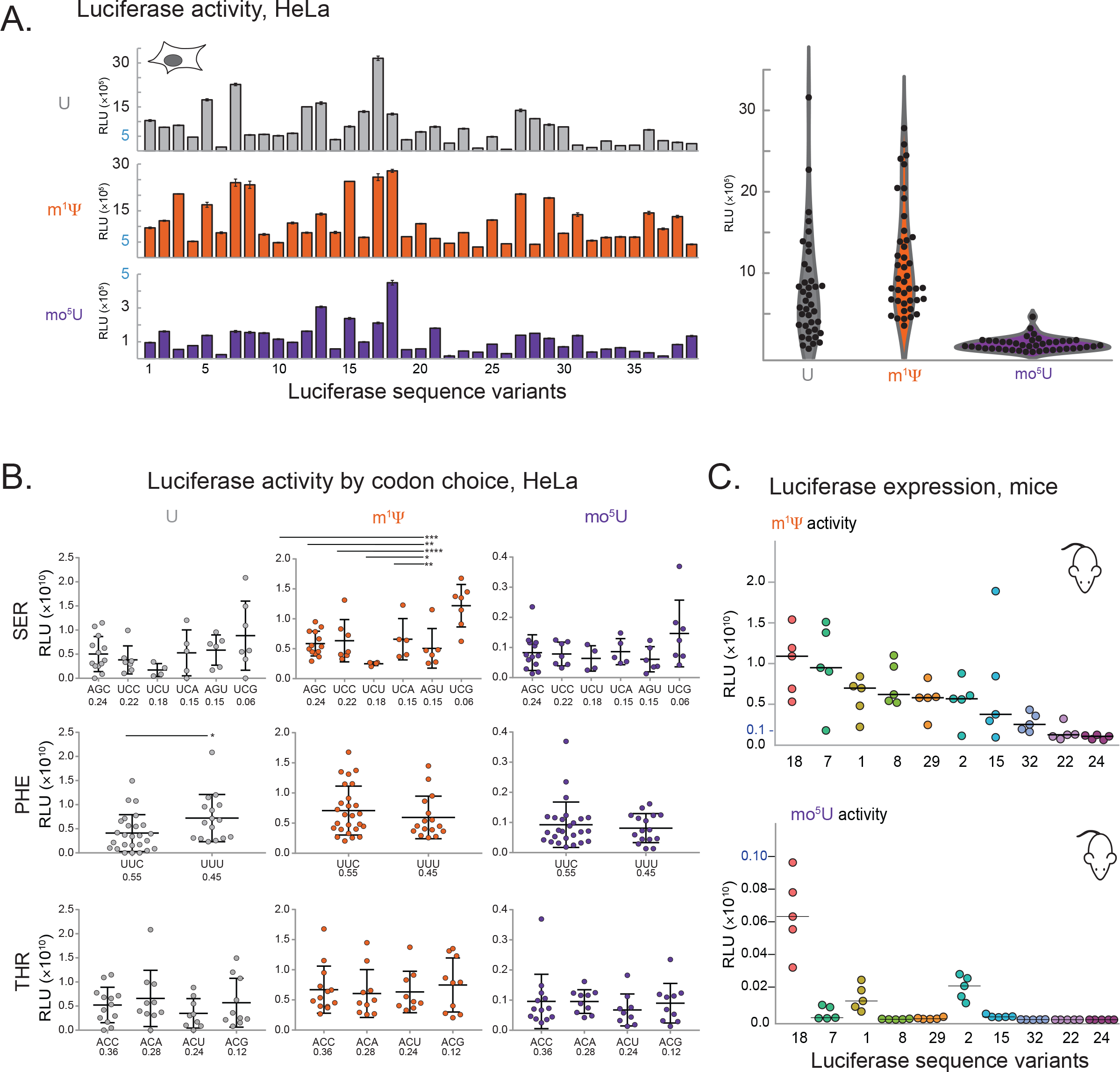
Inclusion of modified nucleotides in mRNA alters Luc expression. (A) Left Panel: expression in HeLa cells (RLU, y-axis) for 39 firefly Luciferase sequence variants (L_1_ through L_39_, x-axis) containing uridine (grey, top), m^1^Ψ (orange, middle), or mo^5^U (dark purple, bottom). Right Panel: distribution of expression levels (RLU, y-axis) for variants (black dots) containing uridine (grey), m^1^Ψ (orange), and mo^5^U (dark purple) as a violin plot. (B) Expression in HeLa cells (RLU, y-axis) of 39 firefly Luciferase variants grouped by the codon used (x-axis) for all instances of serine (top), phenylalanine (middle), and threonine (bottom) in mRNAs containing uridine (left), m^1^Ψ (middle), or mo^5^U (right). Codons are shown in order of frequency of occurrence in the human transcriptome. Individual values (dots) are the same as in A with mean and standard errors (black lines). Significant differences by two-way ANOVA comparisons are indicated by lines above, and *p*-values are noted by asterisks (* *p* ≤ 0.05, ** *p* ≤ 0.01, *** *p* ≤ 0.001, and **** *p*≤ 0.0001). (C) Total luminescence of *in vivo* firefly Luciferase expression (RLU, y-axis) in CD-1 mice (five per group) following IV injection of 0.15 mg/kg LNP-formulated mRNA for 10 sequence variants (x-axis) containing m^1^Ψ (top) or mo^5^U (bottom). Individual animals (dots) with median.

Next, we examined how protein expression was affected by global substitution with modified nucleotides in the same sequences. For eGFP mRNAs in HeLa cells, modified nucleotides changed the expression of both individual variants and the overall expression mean and range of the entire sequence set. Compared to unmodified mRNA, mean expression was similar for eGFP mRNAs containing Ψ and m^1^Ψ, but lower for mo^5^U and Ψ/m^5^C (3-fold and 1.5-fold lower, respectively, Figure 1B). Of note, the identities of the best and worst expressing sequences were not consistent across the different modified nucleotides. For example, eGFP sequence G_2_ expressed highly with Ψ and m^1^Ψ, moderately with U and Ψ/m^5^C, but poorly with mo^5^U (Figure 1B). Similar trends were observed for hEpo mRNA in HeLa cells, with m^1^Ψ yielding a 1.5-fold greater mean expression than U, which was in turn 2-fold higher than mo^5^U (Figure 1C). As with eGFP and Luc, we observed hEpo variants (e.g., E_CO_ and H_A_E_2_) that expressed well with m^1^Ψ, but not U or mo^5^U-containing mRNA (Figure 1C). Although we observed some variation in the expression levels of individual RNAs in hepatocytes versus HeLa cells, the general expression trends were remarkably similar (Figure 1C, Figure S1C).

To extend this analysis, we next examined 39 synonymous Luc sequences containing U, m^1^Ψ, or mo^5^U mRNA in HeLa, AML12 and primary hepatocyte cells. Mean expression increased 1.5-fold for m^1^Ψ mRNA but decreased 5-fold for mo^5^U compared to unmodified mRNA in HeLa cells (Figure 2A). This trend held in AML12 cells and primary hepatocytes cells as well as across delivery methods including electroporation and transfection of lipid nanoparticles (although some individual differences were noted (Figure 2A, Figure S2B, Figure S3B)). For several mRNA sequences, inclusion of modified nucleotides substantially impacted protein expression (Figure 2A, Figure S2C). Several sequences (e.g., L_24_, and L_22_) universally produced low levels of protein across all modified nucleotides, but many variants (e.g., L_18_, L_7_, L_2_, L_8_, and L_29_) favored specific modified nucleotides over others. Taken together, these data indicate that sequence and nucleotide modifications make distinct contributions to the overall level of protein expression.

A simple explanation for the observed modified nucleotide-specific expression differences would be a direct effect on decoding by the ribosome. If so, expression should correlate with overall modified nucleotide content, or alternatively with the use of specific codons containing modified nucleotides. However, there is no clear relationship between % U content and expression (Figure S2A) and only a few m^1^Ψ- and mo^5^U-containing codons had any statistically significant impact on protein output (6 and 4 respectively of 87 synonymous pairwise comparisons *p* < 0.05, Figure 2B and Figure S3A). A notable exception is an unexpected and unexplained 2-fold increase in protein production with inclusion of the non-optimal codon Ser^UCG^ in m^1^Ψ mRNA (*p* < 0.05, Figure 2B and Figure S3A). Thus, codons containing modified nucleotides (Ψ, m^1^Ψ, or mo^5^U) are not inherently translationally deficient.

To assess the degree to which the above conclusions from cell lines translated to animals, we examined protein expression in mice from formulated hEpo and Luc mRNA variants containing two nucleotide modifications shown to have reduced immunogenicity (m^1^Ψ and mo^5^U) (Kariko et al., 2005). Unmodified mRNAs were not included because *in vivo* protein expression can be obscured by strong activation of innate immunity (Kormann et al., 2011). For some hEpo mRNAs, such as m^1^Ψ H_B_E_3_, we noted expression differences between the cell lines and mice (Figure 1C and 1D). These differences were larger than the differences observed between cell lines, and more pronounced for m^1^Ψ hEPO mRNA than for mo^5^U hEPO mRNA (Figure S1D). However, general expression trends were maintained *in vivo* (Figure 1D). All six sequence variants containing m^1^Ψ expressed well (Figure 1D, orange), but only two containing mo^5^U mRNA expressed at detectable levels (Figure 1D, purple). Further, the codon optimized variant E_CO_ expressed well with m^1^Ψ but not at all in mo^5^U. Even so, the best expression came from sequence variants containing mo^5^U (H_A_E_4_ and H_A_E_3_). The mo^5^U H_A_E_4_ variant produced >1.5-fold more protein than the best expressing m^1^Ψ variant (H_A_E_3_, Figure 1D).

The ten Luc variants tested *in vivo* were chosen to represent the widest possible range of protein expression observed in cell culture. As expected from the known biodistribution of MC3-containing LNPs (Sabnis et al., 2018), the liver was the main site of protein expression (Figure S2E). Luc mRNAs containing m^1^Ψ were highly expressed *in vivo*, particularly L_18_ and L_7_ (Figure 2C, top panel).

Variability in protein expression with mo^5^U was more exaggerated *in vivo*, as 7 of the 10 variants produced little to no protein (Figure 2C, bottom panel). L_18_ was an exception, but still produced >10-fold less Luc than the same sequence with m^1^Ψ. Notably, L_7_ produced large amounts of protein with m^1^Ψ but barely detectable levels with mo^5^U (Figure 2C, top versus bottom panel, note the y-axis scales). These data suggest that expression differences observed in cell culture persist and can be more pronounced for exogenous RNAs delivered *in vivo* (Figure S2D).

### Protein expression differences trends with mRNA thermodynamic stability

Since codon usage alone could not fully explain sequence-dependent expression differences in mRNAs containing modified nucleotides, we examined how modified nucleotides might affect mRNA secondary structure. We determined UV absorbance melting curves for mRNAs across a range of expression levels containing different uridine analogs (U, m^1^Ψ, and mo^5^U) as an overall measure of secondary structure. Highly expressing mRNAs underwent substantial melting transitions, detected as sharp peaks in the melting curves, above 35°C (e.g., variant L_18_ with all three uridine analogs and L_15_ with m^1^Ψonly; Figure 3A). For some variants (e.g., L_15_), inclusion of m^1^Ψ but not mo^5^U induced a shift to higher melting temperatures, suggesting global stabilization of structural features within the mRNA (Figure 3A). Notably, L_15_ expression was much higher with m^1^Ψ than mo^5^U (Figure 2A). Similar trends were observed in most, but not all, sequences tested (Figure S4A). Although these initial results suggested a link between RNA structural stability and modification-dependent protein expression *in vivo*, higher resolution structural information was required.

**Figure 3.**
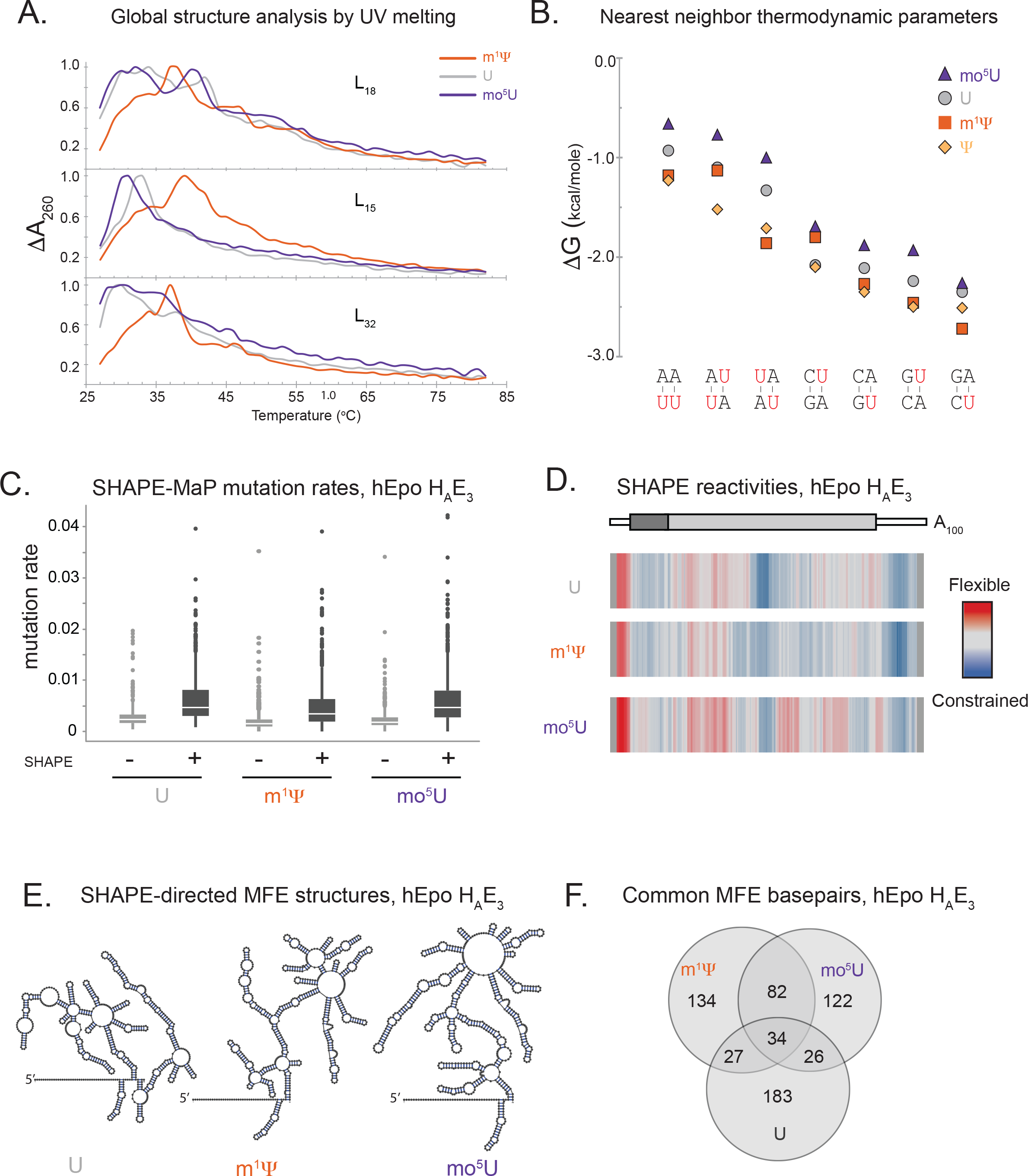
Modified nucleotides induce global structural changes in mRNA. (A) Optical melting profiles of firefly Luciferase sequence variants (L_18_ top, L_15_ middle, and L_32_ bottom) containing uridine (grey), m^1^Ψ (orange), or mo^5^U (dark purple) showing the change in UV absorbance at 260nm (y-axis) as a function of temperature (x-axis). (B) Nearest neighbor thermodynamic parameters for Watson-crick base pairs (x-axis) containing uridine (grey circles, values from (Xia et al., 1998)), Ψ (yellow diamonds), m^1^Ψ (orange squares), or mo^5^U (dark purple triangles). The modified nucleotides in each nearest neighbor are highlighted in red. (C) Apparent mutation rates (y-axis) for untreated (light grey, −) and treated (dark grey, +) samples of hEpo sequence variant H_A_E_3_ containing uridine (grey), m^1^Ψ (orange), or mo^5^U (purple) (x-axis). (D) Median SHAPE reactivity values (33-nt sliding window) for hEpo sequence variant H_A_E_3_ containing uridine (top), m^1^Ψ (middle), or mo^5^U (bottom) shown as a heatmap: highly reactive (red), moderately reactive (grey), and lowly reactive (blue). The positions of the 5’and 3’UTRs (thin white boxes), H_A_ coding sequence (dark grey box), and E_3_ coding sequence (light grey box) are shown in the schematic below. (E) SHAPE-directed Minimum Free Energy (MFE) secondary structure predictions for hEpo sequence variant H_A_E_3_ containing uridine (grey, left), m^1^Ψ (orange, center), and mo^5^U (purple, right). The location of the 5’end of the mRNA is indicated. (F) Venn diagram indicating the number of common and unique base pairs for each SHAPE-directed MFE structures shown in E.

The thermodynamics of base-pairing in RNA is commonly understood in terms of nearest-neighbor energy terms (Lu et al., 2006). Where these parameters were previously reported for unmodified RNA and RNA containing Ψ, to our knowledge they have not yet been established for m^1^Ψ or mo^5^U to our knowledge. To establish these parameters for m^1^Ψ, and mo^5^U, we therefore performed optical melting experiments on 35 synthetic short RNA duplexes containing global substitutions of uridine with Ψ (Xia et al., 1998). Nearest neighbors containing Ψ (Figure 3B, diamonds) and m^1^Ψ (Figure 3B, squares) form substantially more stable base pairs than uridine (by 0.25 and 0.18 kcal/mol on average, respectively; Figure 3B, circles; Table S1). In contrast, nearest neighbors containing mo^5^U (Figure 3B, triangles) are destabilized by 0.28 kcal/mol relative to uridine (Figure 3B, Table S1). The average difference for mo^5^U versus Ψ is −0.5 kcal/mol per nearest neighbor, or −1.0 kcal/mol per base pair. These differences, summed over all base pairs including a modified nucleotide in a full-length mRNA, readily explain the observed differences in the UV melting curves caused by inclusion of different modified nucleotides. Therefore, the rank order of average nearest neighbor base-pairing stability (Ψ ≅ m^1^Ψ > U > mo^5^U, Figure 3B) correlates with mean protein expression (Figures 1 and 2).

### Modified nucleotides induce global rearrangement of mRNA structure

To investigate how modified nucleotides impact mRNA structure at single nucleotide resolution, we used SHAPE-MaP to probe RNA structure (Siegfried et al., 2014). Since the use of SHAPE on RNAs globally substituted with m^1^Ψ and mo^5^U has not been reported, we first validated the methodology. In the absence of the SHAPE reagent (1-methyl-6-nitroisatoic anhydride (1M6)), there was no evidence of increased background error rates by Next-Generation Sequencing (NGS) with either m^1^Ψ or mo^5^U (Figure 3C). However, 1M6 treatment uniformly increased the mutation rates for RNAs containing either m^1^Ψ or mo^5^U to a similar extent as observed for uridine (Figure 3C). Therefore, the SHAPE-MaP method is effective for these globally-modified mRNAs.

Using SHAPE-MaP, we measured RNA structure across the experimentally tested variants of hEpo containing U, m^1^Ψ, or mo^5^U. SHAPE-MaP produces single-nucleotide resolution structural information across entire RNA molecules, with stable structural elements indicated by low SHAPE reactivities and vice versa (Figure S4B, Table S2). Data for a representative sequence, hEpo H_A_E_3_, revealed local structure that differed dramatically by modified nucleotide (Figure 3D). Consistent with the nearest neighbor parameters (Figure 3B) in many RNAs m^1^Ψ stabilized and mo^5^U destabilized structure (hEpo H_A_E_3,_ Figure 3D). These trends are c). Pseudo-free energy constraints based on SHAPE reactivity values were used to model RNA secondary structure with the RNAstructure package (Mathews et al., 2004, Deigan et al., 2009). Such data-informed secondary structure models of hEpo H_A_E_3_ indicate that modified nucleotides induce widespread secondary structure rearrangements even in the context of the same sequence (Figure 3E). The three Minimum-Free Energy models for hEpo H_A_E_3_ containing U, m^1^Ψ, or mo^5^U share fewer than 13% of common base pairs, with the plurality of predicted base pairs being unique to a single modification (Figure 3F). Thus, global incorporation of modified nucleotides induces widespread changes in mRNA structural content and conformation.

### Position-dependent structural context defines highly expressed mRNAs

Having demonstrated that modified nucleotide incorporation induces substantial changes in base-pairing stability and identity, we next investigated the positional dependence of protein expression on local RNA structure. To do so, we obtained SHAPE data for synonymous variants of hEpo (8 variants each with m^1^Ψ or mo^5^U) and Luc (38 variants each with U, m^1^Ψ, or mo^5^U) whose protein output varied over a >2-orders of magnitude (130 mRNAs total). Regions displaying structural differences were identified using 31-nucleotide sliding window median reactivities, as previously described (Watts et al., 2009). Consistent with observations above, high protein output mRNA variants had lower median SHAPE reactivities (i.e., increased structure) across the CDS than low protein output variants. This was true for both proteins and all three chemistries (Figures 4A and 4B; Table S2). Particularly striking examples were E_CO_ and L_8_ mRNAs, where their high expression in m^1^Ψ compared to mo^5^U correlated with widespread m^1^Ψ-dependent decreases in median SHAPE reactivity throughout the CDS (Figures 4A and 4B). In contrast, the 5’ UTR of high expressing variants exhibited high SHAPE, indicating a general lack of structure in this region (Figures 4A and 4B).

**Figure 4.**
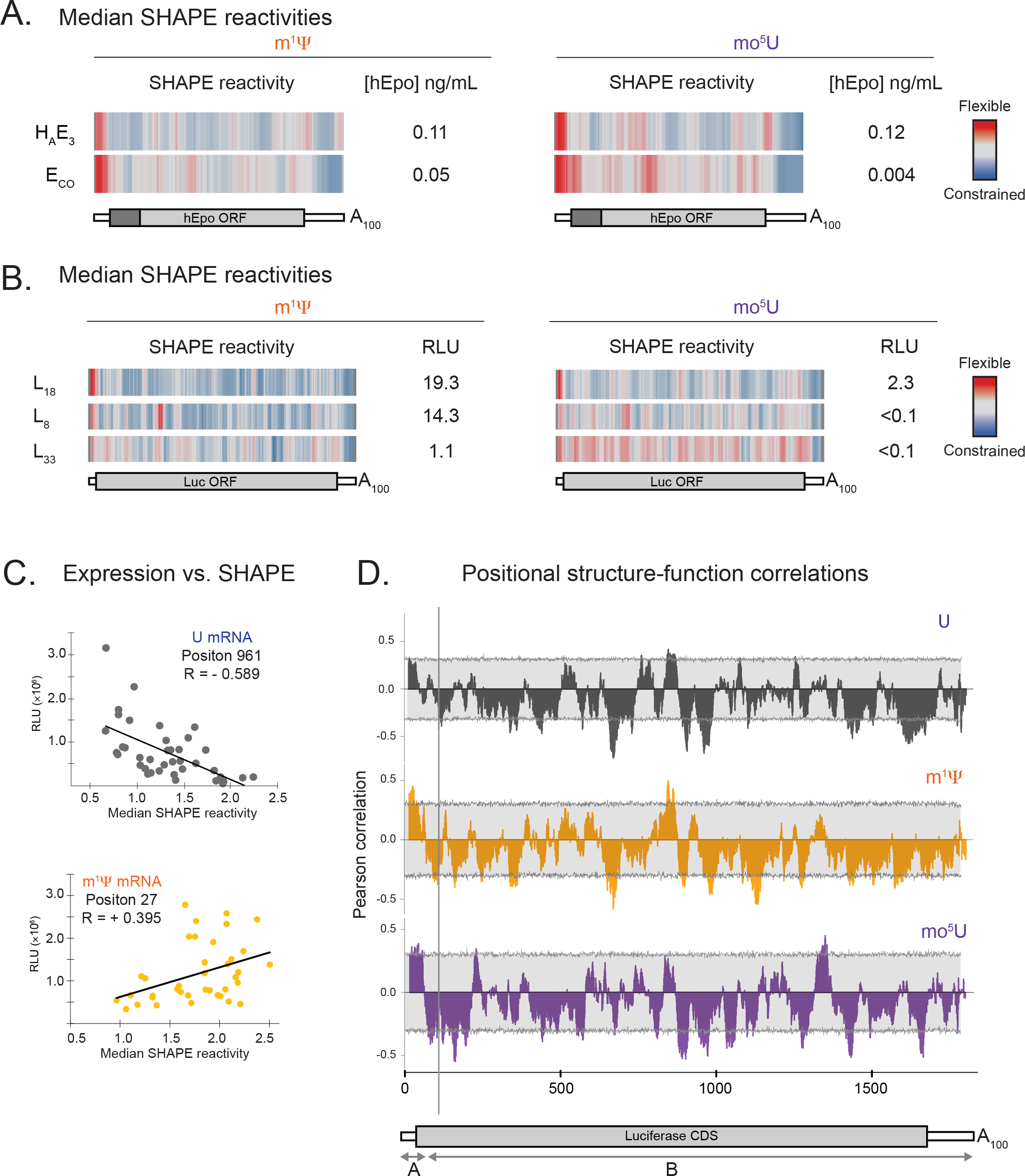
SHAPE data reveal a bipartite relationship between mRNA structure and protein expression. (A) Median SHAPE reactivity values (33-nt sliding window) for hEpo sequence variants E_CO_ (top) and H_A_E_3_ (bottom) containing m^1^Ψ (orange, left) or mo^5^U (purple, right) shown as a heatmap: highly reactive (red), moderately reactive (grey), and lowly reactive (blue). hEpo serum concentrations in mice from Figure 1D are shown to the right. The positions of the 5’and 3’UTRs (thin white boxes), H_A_ coding sequence (dark grey box), E_2_ coding sequence (light grey box), and poly A tail are shown in the schematics below. (B) As in A above but for firefly Luc sequence variants L_18_ (top), L_8_ (middle), and L_32_ (bottom). Total luminescence values in mice from Figure 2C are shown to the right. (C) Expression in HeLa cells (y-axis, from Figure 2A) for 39 firefly Luciferase variants (dots) containing uridine (dark grey, top) or m^1^Ψ (orange, bottom) versus median windowed SHAPE reactivity value (x-axis) in two 33-nt windows centered at the indicated positions. (D) Pearson correlation (y-axis) versus nucleotide position (x-axis) for firefly 39 Luciferase sequence variants containing U (dark grey, top), m^1^Ψ (orange, middle) or mo^5^U (dark purple, bottom). The light grey boxes show the empirical 95% confidence interval at each position.

We next analyzed the directionality and strength of the correlation between position-wise SHAPE reactivity and protein expression across all Luc variants (Figure 4C). This revealed a striking, position-dependent relationship between mRNA structure and expression that was largely consistent between mRNAs with m^1^Ψ and mo^5^U. The region encompassing the 47-nt 5’ UTR and the first ~30 nucleotides of the CDS (Figure 4D; region A) showed a positive and statistically significant correlation (*p*< 0.05) between SHAPE reactivity and protein expression for both m^1^Ψ and mo^5^U mRNAs. In contrast, the remainder of the CDS and the entire 3ʹ UTR (Figure 4C, region B) exhibited a predominantly inverse correlation between SHAPE reactivity and protein expression for U, m^1^Ψ, and mo^5^U. In other words, increased secondary structure in these regions correlated with improved protein expression, consistent with the global structural properties measured by optical melting. Notably, however, the strength of the structure-function correlation varied across the meta-sequence, with specific regions exhibiting statistically-significant negative (e.g., around position 961) or positive (e.g., around position 850) correlations between SHAPE reactivity and protein expression.

The above analyses suggest that structural changes induced by modified nucleotides directly impact protein expression. We examined the role of flexibility in region A (47-nt 5’ UTR and the first 30 nucleotides of the CDS) by creating chimeras combining variants with different structural signatures. Luc variants L_7_ and L_27_ (Figure 2A) both exhibited lower than average protein expression in m^1^Ψ. Both also exhibited low SHAPE reactivity (high structure) throughout both regions A and B (Table S2). However, when we replaced region A with the relatively unstructured corresponding region A from the high expresser L_18_ (Figure 4B) to produce fusion mRNAs FL_18/7_ and FL_18/27_, both region A SHAPE reactivity and Luc expression increased (Figure 5A and 5B). The FL_18/7_ and FL_18/27_ chimeras only differed by 2 and 4 individual bases from their respective parents (note that the 47-nucleotide 5’ UTR is common to all sequences). Thus, relatively small changes in the first 30 nucleotides of the coding sequence that drive structural rearrangements can have substantial impacts on protein output.

**Figure 5.**
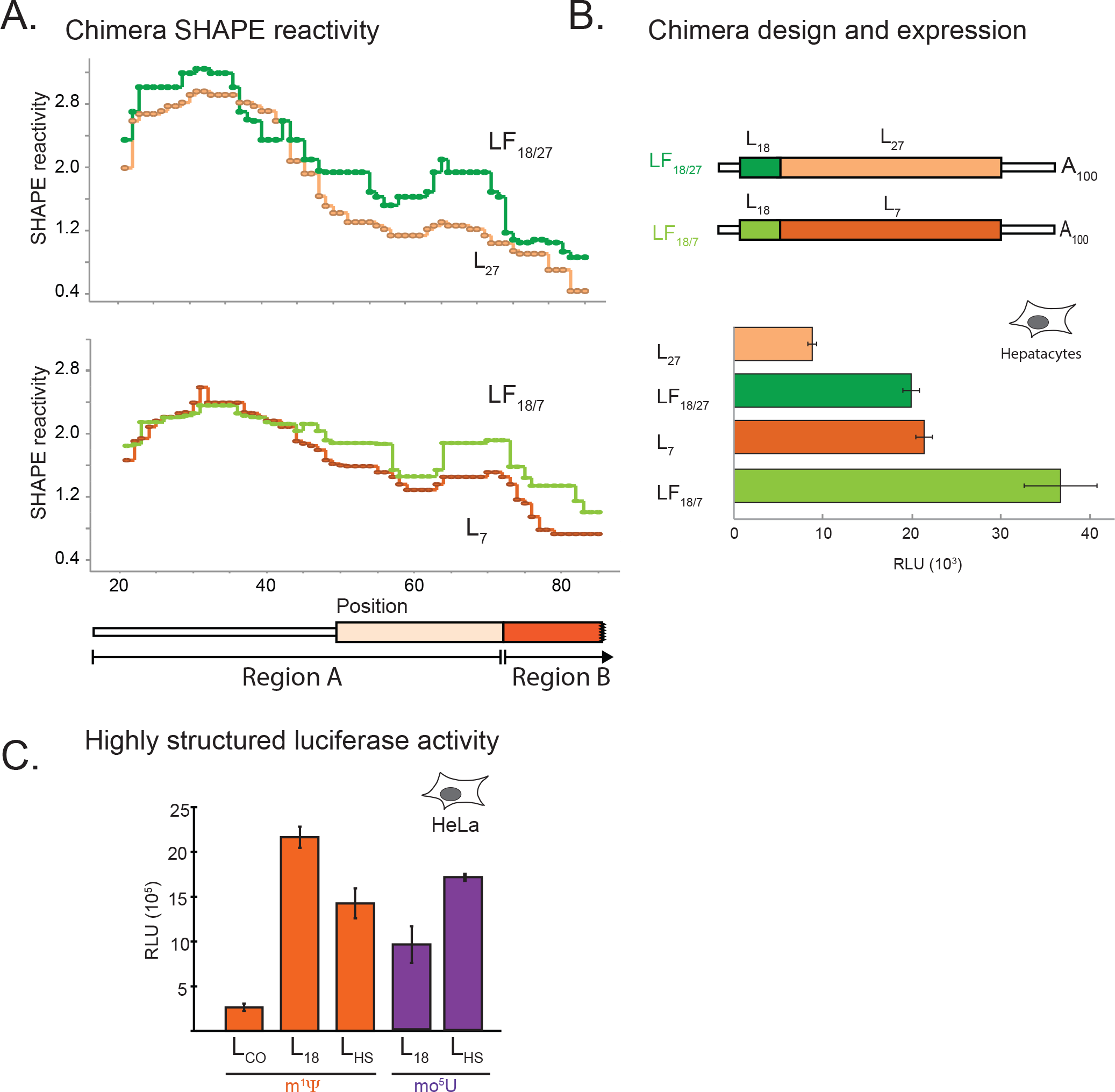
RNA Structure and codon usage combine to determine protein expression. (A) Median SHAPE reactivity values (y-axis, 33-nt sliding window) for firefly Luciferase sequences containing m^1^Ψ (top: LF_18/27_ dark green and L_27_ peach, bottom: L_18/7_ light green and L_7_ red) versus nucleotide position (x-axis) for the first 100 nucleotides. The positions of the 5’UTR (thin white box), the beginning of the CDS (colored boxes) are shown. Lines indicate the position of the luciferase region swapped in the chimeric constructs (Region A) (B) Top: schematic of 2 chimeric constructs (LF_18/27_ top and LF_18/7_ bottom) which combine 77 nt regions near the start codon with the remainder of the CDS of different firefly Luciferase sequence variants. Bottom: expression in primary mouse hepatocytes (RLU, x-axis) for original (L_7_ and L_27_) and fusion (LF_18/27_ and LF_18/7_) firefly Luciferase constructs (y-axis) containing m^1^Ψ. (C) Expression in HeLa cells (RLU, y-axis) for firefly Luc sequence variants from Figure 2A (L_18_, L_CO_) or engineered to have more stable secondary structure (L_HS_) containing m^1^Ψ (orange) and mo^5^U (dark purple).

To investigate the degree to which secondary structure within region B (rest of the CDS and the 3’ UTR) drives protein output, we designed a Luciferase variant (L_HS_) in which the coding sequence was computationally predicted to have more stable secondary structure in region B than any of our previously tested luciferase mRNAs (L_HS_ for High Structure). This high degree of region B structure was confirmed by SHAPE reactivity (Supplemental Table S2). In the mo^5^U chemistry, the L_HS_ variant yielded 1.5-fold greater protein expression than L_18_, the previous best expressing variant in mo^5^U (Figure 5C). For mRNAs containing m^1^Ψ, L_HS_ expression of reduced compared to L_18_ and was also slightly below L_HS_ RNA containing mo^5^U. Thus, both removing structure in region A or increasing structure in region B through sequence alterations can dramatically impact protein expression.

### Codon usage and mRNA structure synergize to determine ribosome loading and mRNA half-life

The redundancy of the genetic code means that for an average sized mammalian protein (~430 aa; (Scherer and Cold Spring Harbor Laboratory. Press., 2010)) there are >10^150^ synonymous coding sequences. Therefore, it is impossible to completely enumerate the relationships between codon usage, mRNA secondary structure and protein expression. Instead we computationally generated sets of 150,000 synonymous CDSs encoding eGFP-degron (284 amino acids with 2.3 ×10^135^ possible synonymous mRNA sequences) with 3 different algorithms. For each sequence, we calculated Relative Synonymous Codon Usage (RSCU, (Scherer and Cold Spring Harbor Laboratory. Press., 2010)) and the predicted MFE structure (Lu et al., 2006). Randomly choosing synonymous codons with equal probability generates sequences that cluster around 0.75 +/− 0.05 RSCU and −325 +/− 40 kcal/mol MFE (Figure 6A, red). Using probabilities weighted by frequency in the human transcriptome (Nakamura et al., 2000) generates a similar-shaped distribution, but shifted to both significantly higher RSCU (0.825 +/− 0.05, *p* < 0.05) and greater structure (−340 +/− 40 kcal/mol, *p* < 0.05) (Figure 6A, green). Next, we developed an algorithm that varied the probabilities of individual codon choices dynamically so that RCSU and MFE were both driven to their accessible extremes (Figure 6A, grey). The space covered is far greater than for random or frequency-weighted sequences, but has well-defined limits. Notably, because optimal codons tend to be GC-rich the structure of the genetic code inherently disallows sequences with both highly optimal codons and low structure (top left corner) or rare codons and high structure (bottom right corner).

**Figure 6.**
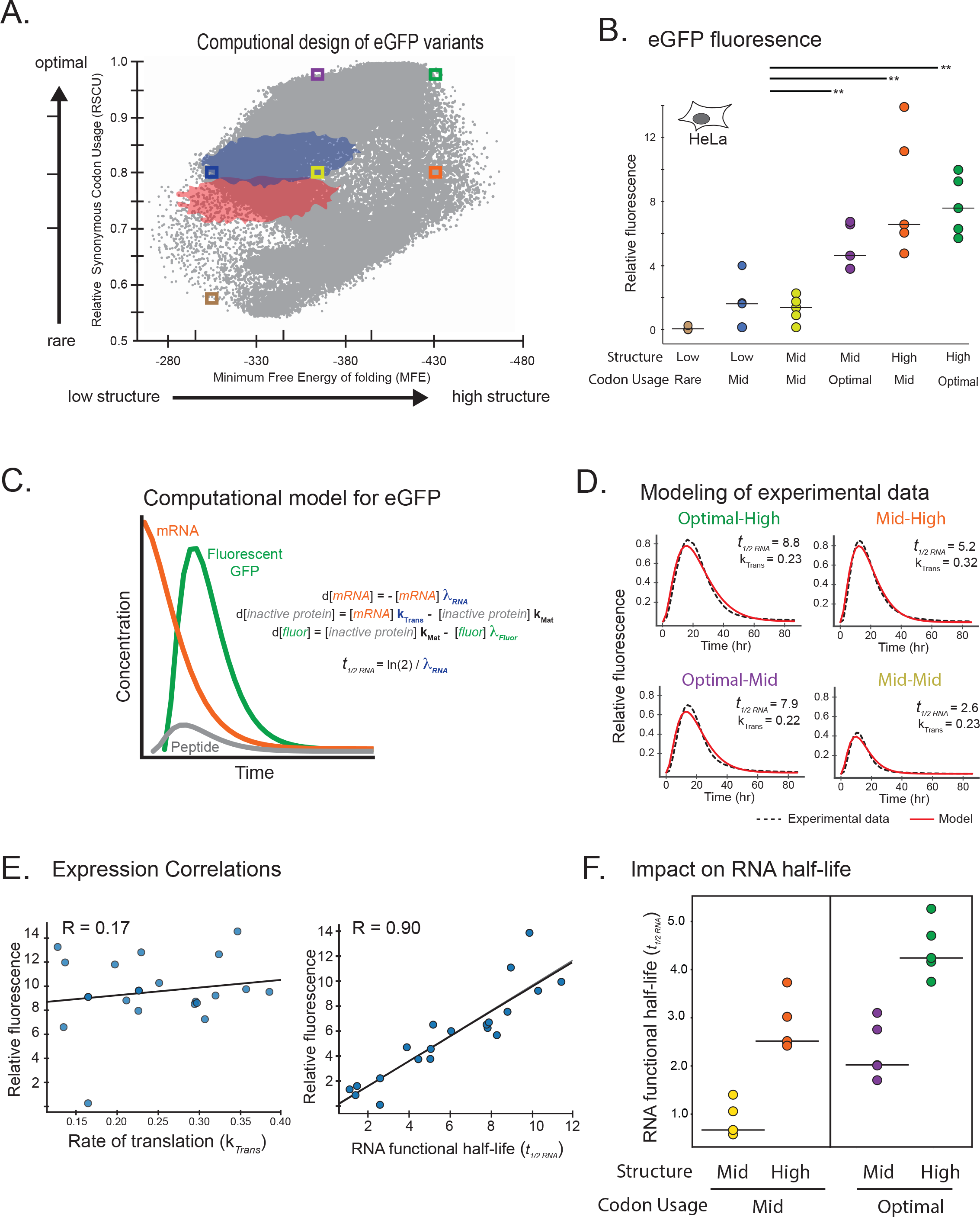
Half-life of computationally designed eGFP-degron mRNAs is determined by codon usage and mRNA structure. (A) Codon optimality (Relative Synonymous Codon Usage, y-axis) versus secondary structure (energy of the predicted MFE structure, x-axis) for sets of 150,000 generated eGFP sequence variants generated using codons chosen randomly (red), weighted in proportion to the human genome(blue), and using our algorithm(grey). Colored boxes show regions from which sequences were selected for further testing. (B) Total integrated eGFP fluorescence measured every 2 hours for 86 hours in HeLa cells (RFU, y-axis) for six sets of five mRNAs containing m^1^Ψ (dots, with median as black line) with differing degrees of codon optimality and/or secondary structure (x-axis, as in A). Significant differences by two-way ANOVA comparisons are indicated by lines above, and *p*-values are noted by asterisks (** *p* ≤ 0.01). (C) Model of eGFP expression kinetics. Simulated curves based on equations for changes in levels of mRNA (mRNA, orange), immature non-fluorescent protein (inactive protein, grey), and mature fluorescent protein (fluor, green) over time using exponential decay rates for mRNA (λ_RNA_) and eGFP protein (λ_Fluor_), and rates of translation (k_Trans_) and protein maturation (k_MAT_). mRNA half-lives (*t_1/2 RNA_*) were calculated from the observed mRNA decay rates. (D) eGFP-degron fluorescence in HeLa cells (RFU, y-axis) versus time (x-axis) as measured experimentally (solid colored lines as in A) and fitted according to the model in C (dashed black lines) for representative sequence variants with differing degrees of codon optimality and/or secondary structure (as in A). Translation rate constants (k_Trans_) and mRNA half-lives (*t_1/2 RNA_*) as derived from the model described in C are shown. (E) Total eGFP-degron fluorescence in HeLa cells (RFU, y-axis) versus the modeled rate constants for translation (k_Trans_, left) or mRNA functional half-life (λ_RNA_, right) for 20 sequence variants containing m^1^Ψas in Figure 5D. Linear regression (black line) and Pearson correlation are shown. (F) Modeled functional mRNA half-lives (λ_RNA_, y-axis) for four sets of five eGFP-degron sequence variants with differing degrees of codon optimality and/or secondary structure (x-axis, as in A and B).

To investigate how the limits of codon optimality and allowable structure affect protein expression, we selected six regions collectively spanning the range of accessible space (Figure 6A, boxes). From each of these six selected regions, we synthesized five synonymous sequences (30 in total) and followed the production and decay of GFP fluorescence over a 20-hour timecourse in HeLa cells. This enabled us to directly compare the effects changing each factor independently, for example changes in MFE at constant RSCU (Figure 6A, yellow vs orange or purple vs green) or the converse (Figure 6A, yellow vs purple or orange vs green). As expected, eGFP-degron mRNAs containing rare codons and very little secondary structure produced minimal protein (Figure 6B, brown). Low protein expression was also observed for mRNAs with middling scores in both relative synonymous codon usage and structure (Figure 6B, yellow). Notably, increasing either the codon optimality or secondary structure while holding the other feature constant both increased median protein expression (Figure 6B, orange and purple respectively, *p* value < 0.01). The set of mRNAs with the highest codon optimality and most structure, however, showed no additional increase in median protein expression (Figure 6B, green, *p* value = 0.55). Similar effects were observed in AML12 cells (Figure S5). Combined, these data indicate that codon usage and secondary structure are both important, but distinct, regulators of overall protein expression.

Next, we analyzed the kinetics of protein production. Real-time, continuous expression data were fit by a model including rate constants for mRNA translation, mRNA functional half-life, maturation of eGFP protein into its fluorescent form (Crameri et al., 1996), and eGFP protein degradation (Figure 6C). Functional half-life reflects the productive life of the mRNA in generating protein and is not necessarily the same as physical half-life ending with degradation – it could also reflect intracellular trafficking or sequestration away from the ribosomal machinery. Since all mRNA sequences expressed the same eGFP protein sequence and we measured fluorescent (i.e., mature functional) protein, we could assume constant rates of protein maturation (k_Mat_) and protein degradation (λ_Fluor_, Figure 6C). Fitting this model to the experimental data allowed us to calculate the rate of translation (k_Trans_) and functional half-life (t_1/2 RNA_) individually for each mRNA variant (Figure 6D). Surprisingly, whereas overall protein expression correlated poorly with mRNA translation rate (*r* = 0.17), it correlated remarkably well with functional mRNA half-life (*r* = 0.90, Figure 6E). These results were consistent across multiple computational models including models containing a delivery rate (Figure S5B and C). Highly structured mRNAs had a >2-fold increase in functional mRNA half-life relative to those with middling degrees of secondary structure, regardless of whether their codon usage was middling or optimal (Figure 6F). Thus, secondary structure increases protein output by extending mRNA functional half-life in a previously unrecognized regulatory mechanism independent of codon optimality.

## Discussion

The amount of protein produced from any given mRNA (i.e., the translational output) is influenced by multiple factors specified by the primary nucleotide sequence. These factors include GC content, codon usage, codon pairs, and secondary structure. Disentangling the individual roles played by each of these factors in translational output of endogenous mRNAs, however, has proven difficult because of their high covariance. For example, optimal codons tend to be GC heavy, and high GC content drives secondary structure. To separate these confounding relationships, we directly manipulated the secondary structure of exogenously-delivered mRNAs using two distinct approaches. First, we globally replaced uridine with modified versions having markedly different base-pairing thermodynamics – this led to global secondary structure changes without altering the mRNA sequence. Second, we used computational design to identify sets of mRNAs whose coding sequences explored the very limits of codon usage and secondary structure.

Global incorporation of different modified nucleotides often (but not always) markedly changed mRNA expression. This effect was seen across numerous synonymous coding variants of multiple proteins, in several different cell lines, and *in vivo* (Figures 1 and 2). Although there were exceptions, m^1^Ψ generally gave higher expression than U or mo^5^U for the same sequence. Biophysical studies revealed that m^1^Ψ and mo^5^U have dramatically different and opposite effects compared to U (stabilizing and destabilizing, respectively) on overall mRNA folding, nearest-neighbor base pairing thermodynamics, and secondary structure pattern as mapped by SHAPE (Figure 3). We also found that secondary structure correlates with protein expression in a position-specific manner (Figure 4). In highly expressed mRNAs, the entire 5’ UTR and the first ~30 CDS nucleotides generally had low structure content. Notably, even though the constant, 47-nucleotide 5’ UTR was chosen to support high expression across many CDS and we still observed a clear structure-expression relationship in this region. This is consistent with previous reports that a relatively unstructured 5'-end is associated with higher protein expression (Ding et al., 2014; Gu et al., 2010; Shah et al., 2013; Tuller and Zur, 2015; Wan et al., 2014), and with our current understanding of translation initiation, which requires accessibility of the cap and region around the start codon to the canonical translation initiation factors and small ribosomal subunit. Unexpectedly, however, we found that a highly structured CDS region also correlates with increased protein expression. By rationally designing sequences to contain more CDS structure, we could rescue low expressing mo^5^U-containing mRNAs (Figure 5). Protein expression from sequences selected to vary the degree of secondary structure independent of codon usage, and *vice versa*, revealed that secondary structure and codon usage each have distinct and roughly equivalent impacts on protein output (Figure 6). For this set of mRNAs, total protein output was driven primarily by changes in functional mRNA half-life.

What is the mechanistic basis for the observed relationship between CDS secondary structure and functional mRNA half-life? Numerous RNA binding proteins (RBPs) interact with secondary structures, with the RNA helicase DDX6 and double-stranded RNA binding protein Staufen both having been reported to positively impact translation (Jungfleisch et al., 2017). It is also possible that more structured RNAs are less prone to cleavage by single strand-specific endonucleases, although endonucleolytic cleavage is thought to only contribute to the degradation in specific cases (Schoenberg, 2011; Wilamowski et al., 2018). A third possibility is that more structure leads to slower ribosomes, and slower ribosomes are beneficial for enhanced functional protein output and extended mRNA half-lives. For many mRNAs encoding complex multi-domain proteins, ribosome pausing at specific locations is crucial for proper protein folding, and therefore activity (Kimchi-Sarfaty et al., 2007; Rauscher and Ignatova, 2018). Although most such pauses are driven by selective maintenance of rare codons (Buhr et al., 2016), regions of high secondary structure may play a similar role in HIV mRNAs encoding proteins with inter-domain junctions (Watts et al., 2009). This latter hypothesis is consistent with biophysical experiments clearly demonstrating an effect of secondary structure on slowing ribosome processivity (Wen et al., 2008).

But how might slower ribosomes lead to an increase in the number of productive rounds of translation? Ribosomal pauses induced by rare codons are directly linked to mRNA degradation (Presnyak et al., 2015). However, the impact and frequency of ribosomal pausing induced by secondary structure is not well understood. Although secondary structure likely slows ribosome progression, the ribosome is inherently a very powerful helicase that necessarily unwinds mRNA structure during elongation (Mustoe et al., 2018). Local secondary structure elements unwound by each advancing ribosome should quickly reform after that ribosome has moved on, and may thus act as “buffers” to prevent collisions between adjacent ribosomes on the same message. Notably, ribosome collisions were recently demonstrated to activate the No-Go mRNA decay (NGD) pathway and decrease mRNA half-life (Simms et al., 2017). Even more recent work has shown that small subunit proteins of collided di-ribosomes are targets of ZNF598 ubiquitin ligase (Juszkiewicz et al., 2018). Which could be a crucial marking event targeting both the ribosome and it bound mRNA for degradation. So, the explanation for how more local secondary structure extends functional mRNA half-life may simply be a protective buffering effect that increases the spacing between ribosomes and thereby decreases collisions (Zarai et al., 2016).

## Author Contributions

Conceptualization, D.M.M, B.J.C, I.J.M., J.R., V.P, D.R., M.M; Investigation, D.M.M, B.J.C, V.P, D.W.R, N.K, S.V.S, B.G, K.L ; Formal Analysis, D.M.M, B.J.C, V.P., D.W.R, N.K, S.V.S, B.G, K.L; Writing – Original Draft, D.M.M and B.J.C; Writing – Review & Editing, D.M.M, B.J.C, V.P., D.W.R., S.V.S., I.J.M., M.J.M ; Visualization, D.M.M, B.J.C, I.J.M., V.P, M.J.M ; Supervision, D.M.M, K.L, D.R., V.P, S.V.S, J.R., I.J.M., M.J.M

## Acknowledgements

We would like to thank DNA Software for expert work in the experimental determination of the nearest neighbor parameters for modified nucleotides, Moderna’s Preclinical Production group for production and formulation of mRNA, Moderna’s In Vitro Biology group for cell culture experiments, Moderna’s Nonclinical Sciences group for *in vivo* experiments, Chris Pepin and Wei Zheng for statistics advice and support, and Paul Yourik, Mihir Metkar, Alicia Bicknell, Ruchi Jain, and Caroline Köhrer for critical reading of the manuscript. This work was supported by Moderna Therapeutics.

## Conflict of interest statement

All authors are employees (or ex-employees in the case of SS, J.R, and J.C.) of Moderna Therapeutics.

## Supplemental Figure/Table Legends

**Figure S1.**
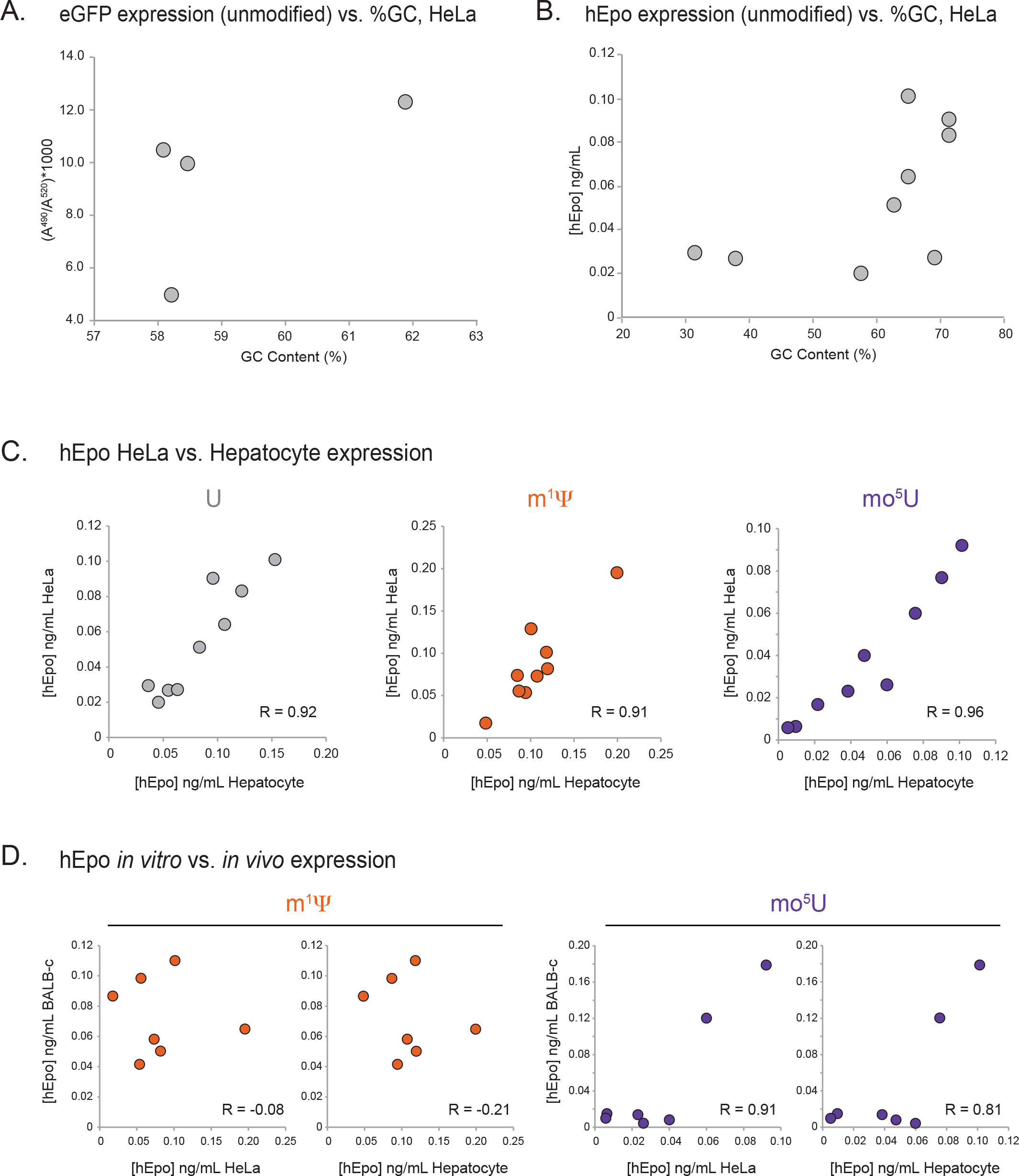
Inclusion of modified nucleotides in mRNA alters protein expression, related to Figure 1. (A) Correlation between the %U (left panels), GC% (middle panels), and RSCU (right panels) for mRNA (x-axis) and eGFP protein production in HeLa cells (y-axis) for mRNA containing uridine (top panel), m^1^Ψ (middle panels), or mo^5^U (bottom panels). (B) Distribution of expression levels (RLU, y-axis) for in AML12 (top panel) and primary mouse hepatocytes (bottom panel) for mRNA variants containing uridine (grey), m^1^Ψ (orange), and mo^5^U (dark purple) shown as a violin plot. (C) Correlation of secreted hEpo protein production in primary mouse hepatocytes (x-axis) and HeLa cells (y-axis) as measured by ELISA in ng/mL following transfection of cells with 8 sequence variants (described in B above) plus one “codon optimized” variant (E_CO_) (Welch et al., 2009) containing uridine (left panel), m^1^Ψ (middle panel), or mo^5^U (right panel). (D) Correlation of secreted hEpo protein production in primary mouse HeLa cells (right graph) and primary mouse hepatocytes (left graph) to mean serum concentrations (y-axis) of hEpo protein in BALB-c mice following IV injection of LNP-formulated mRNA of 6 sequence variants plus one “codon optimized” variant (E_CO_) (Welch et al., 2009). Data is shown for mRNA containing m^1^Ψ (left panel) and mo^5^U (right panel). (E) m^1^Ψ (middle panel), or mo^5^U (right panel)

**Figure S2.**
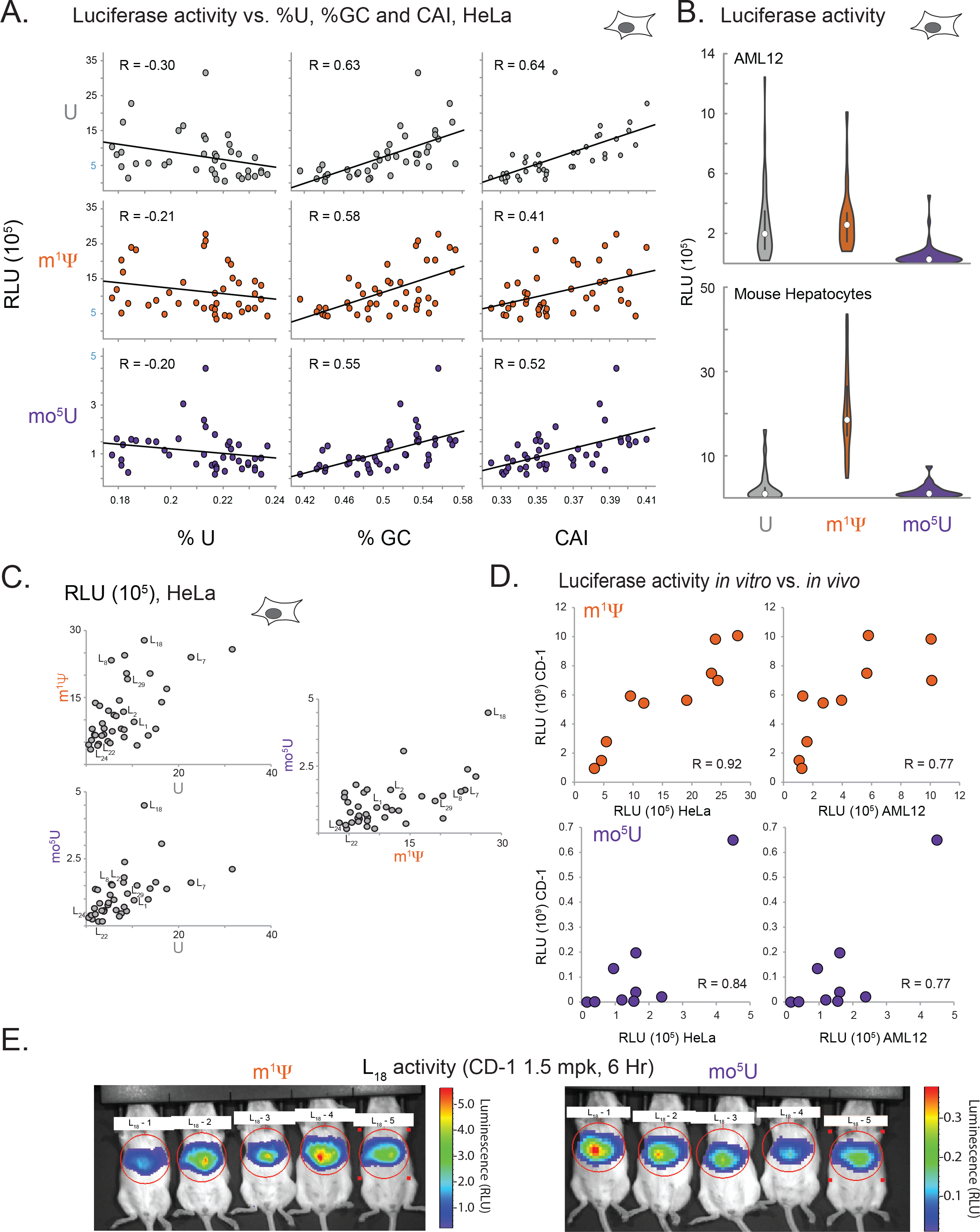
Inclusion of modified nucleotides in mRNA alters Luc expression, related to Figure 2. (A) Correlations between U% (x-axis, left column), GC% (x-axis, middle column), or codon adaptive index (CAI) (x-axis, right column) vs. Luc expression in HeLa cells (RLU) (y-axis) for 39 Luc sequence variants containing U (top row), m^1^Ψ (middle row), and mo^5^U (bottom row), with linear regressions and Pearson correlations. Values are the same as in Figure 2A. (B) The distribution of expression levels across all variants for each nucleotide is shown as a violin plot with the median (white circle) and inter-quartile range (black lines) of expression values indicated for uridine (grey), m^1^Ψ (orange), and mo^5^U (dark purple). Distribution shown for expression levels in both AML12 cells (top panel) and primary mouse hepatocytes (bottom panel). (C) *In vivo* Luc expression as measured for a group of five mice injected with formulated L_18_ mRNA containing either m^1^Ψ (left) or mo^5^U (right). Heatmap of luminescence values are shown for each panel along with pixels (red circle) used to quantify total luminescence in Figure 2E. (D) Correlation of Luc protein production in primary mouse HeLa (right graph) and AML12 (left graph) cells to mean total luminescence of *in vivo* protein expression (RLU, y-axis) in CD-1 following IV injection of 1.5 mg/kg LNP-formulated mRNA for 10 Luc sequence variants containing m^1^Ψ (left panel) or mo^5^U (right panel). (E) Whole body luminescence of CD-1 mice (five per group) following IV injection of 0.15 mg/kg LNP-formulated mRNA for 10 sequence variants (x-axis) containing m^1^Ψ (left panel) or mo^5^U (right panel). Luminescence in the circled regions were used to quantify total expression shown in figure 2C.

**Figure S3.**
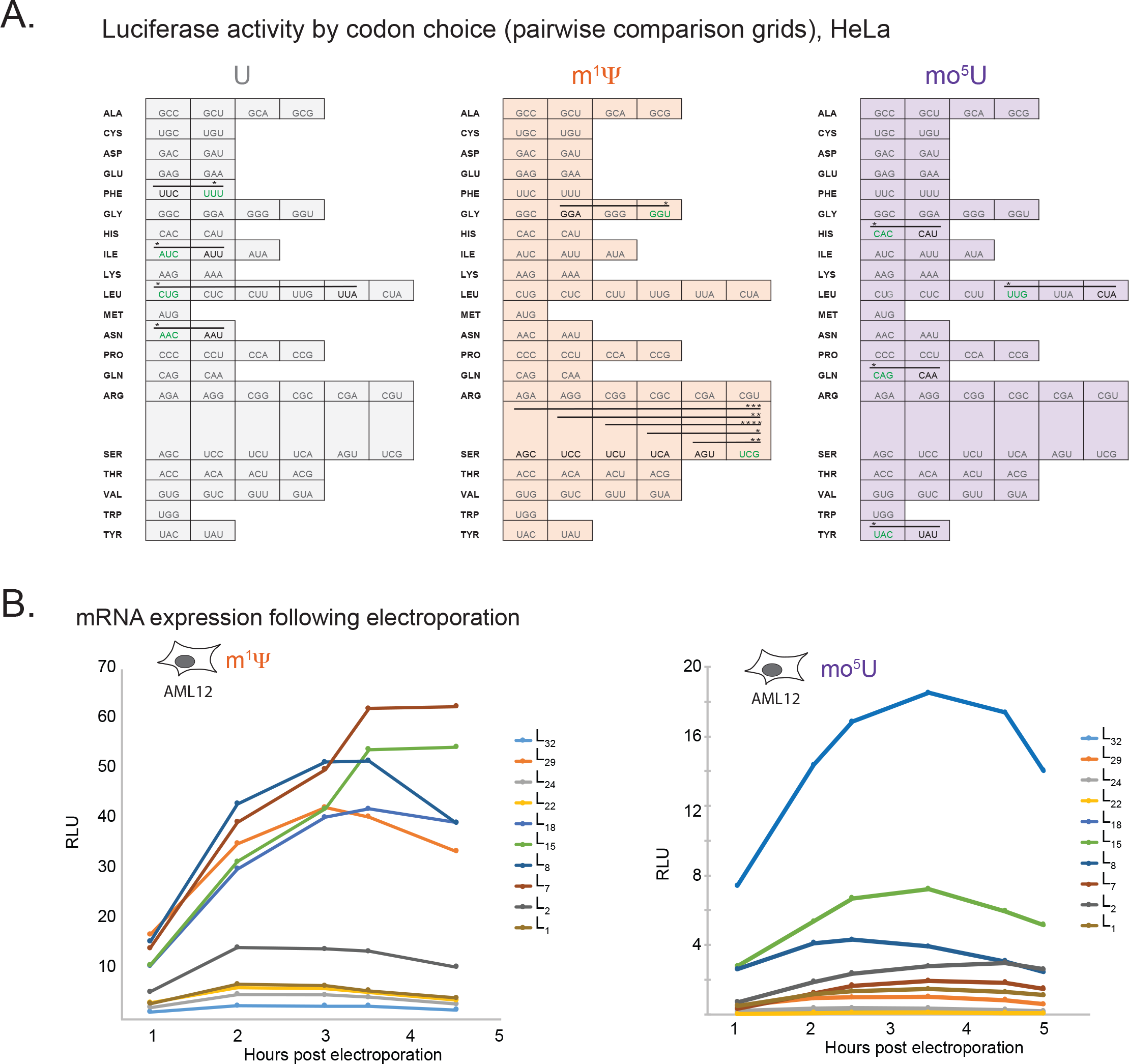
Codon effects of inclusion of modified nucleotides on Luc expression, related to Figure 2. (A) Grid comparisons of protein expression for 39 Luc sequence variants by global codon usage (rows) for mRNA containing uridine (left grid), m^1^Ψ (middle grid), or mo^5^U (right grid). Each row is ordered by frequency of codons in human genome with the most frequent appearing on the left. Codons for which global usage does not significantly impact protein expression relative to other codons are colored grey. Significant differences by two-way ANOVA comparisons are indicated using lines and the codon with the higher median expression value is colored green. P-values are noted by an increasing number of asterisks for P ≤ 0.05 (*), ≤0.01 (**), ≤0.001 (***), and ≤0.0001 (****). (B) Expression in HeLa cells (RLU, y-axis) for 12 firefly Luc sequence variants(x-axis) following electroporation of mRNA containing m^1^Ψ (left panel), or mo^5^U (right panel).

**Figure S4.**
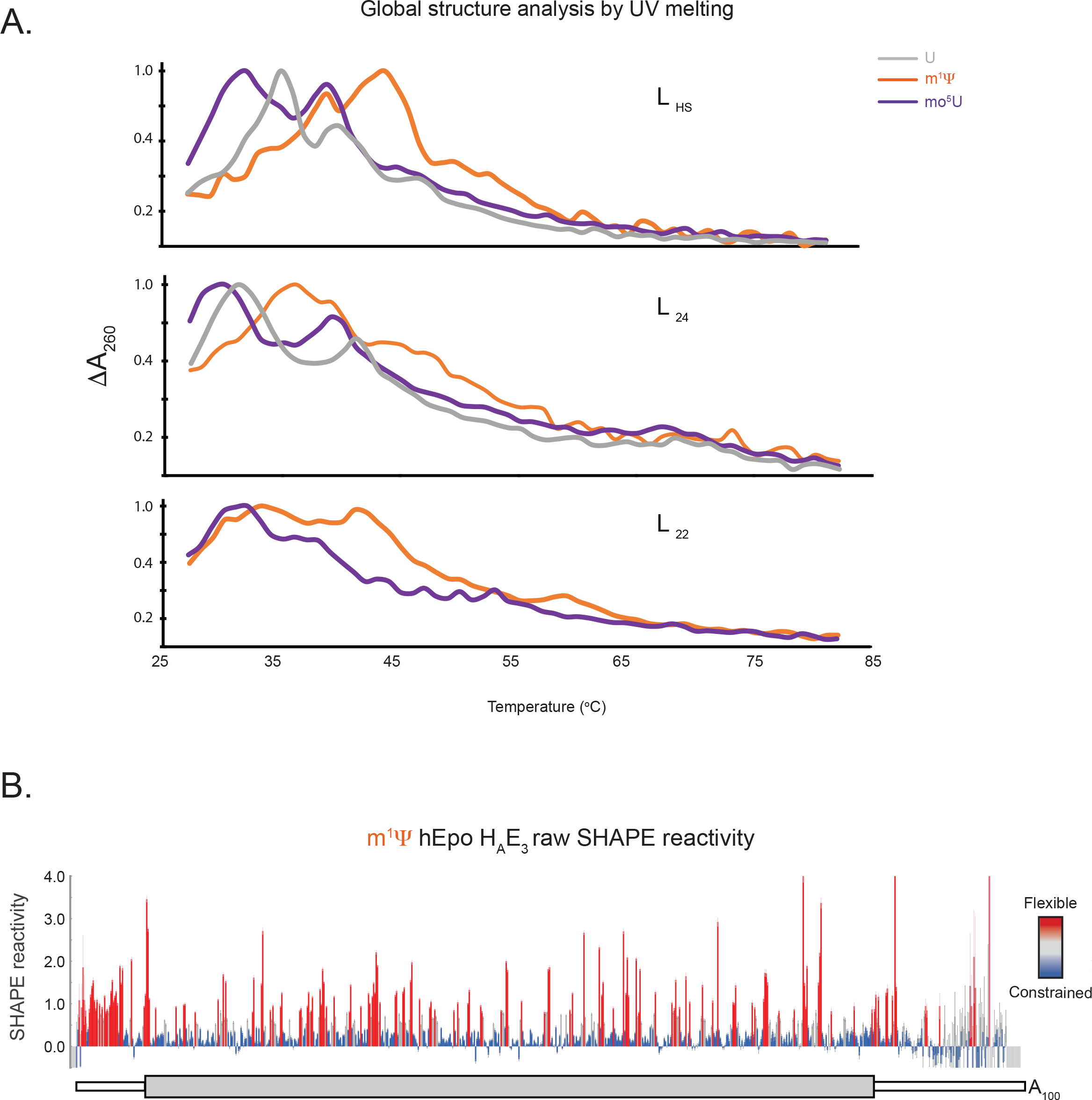
Optical melting of hEPO mRNAs, related to Figure 3. (A) Optical melting profiles of Luc sequence variants L_18_, L_15_, and L_32_ containing uridine (grey), m^1^Ψ (orange), or mo^5^U (dark purple) showing the change in UV absorbance at 260nm (y-axis) as a function of temperature (x-axis). (B) SHAPE reactivity values for each nucleotide in hEpo sequence variant H_A_E_3_ containing m^1^Ψ shown as a column graph with errors. Colored columns indicate highly reactive (red), moderately reactive (grey), and lowly reactive (blue) nucleotides. The positions of the 5’ and 3’ UTRs (thin white boxes), H_A_ coding sequence (dark grey box), and E_3_ coding sequence (light grey box) are shown in the schematic below.

**Figure S5.**
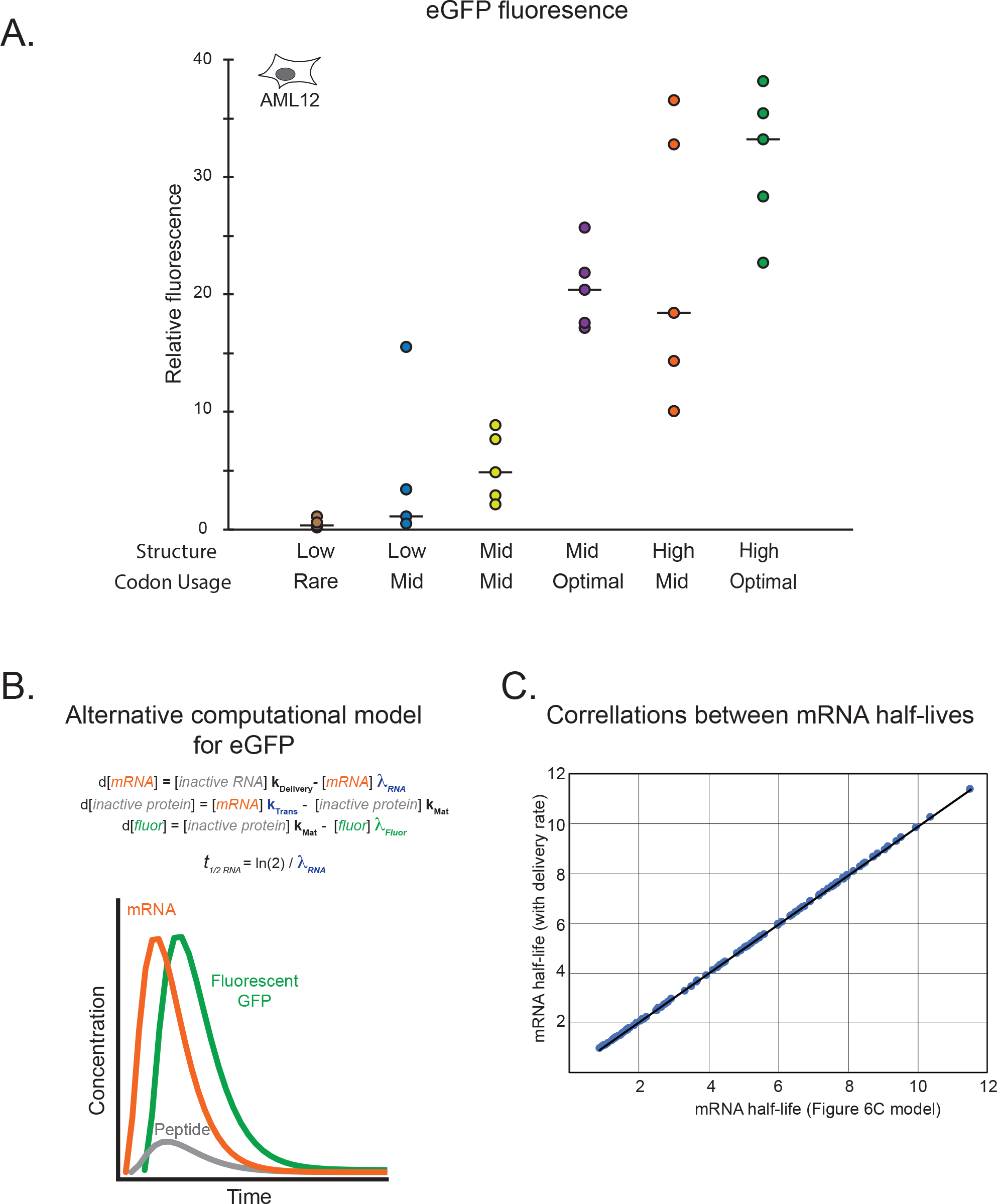
Expression of designed eGFP mRNAs, related to Figure 6. (A) Total integrated eGFP fluorescence measured every 2 hours for 86 hours in AML12 cells (RFU, y-axis) for six sets of five mRNAs containing m^1^Ψ (dots, with median as black line) with differing degrees of codon optimality and/or secondary structure (x-axis, as in A). (B) Model of eGFP expression kinetics. Simulated curves based on equations for changes in levels of mRNA (mRNA, orange), immature non-fluorescent protein (inactive protein, grey), and mature fluorescent protein (fluor, green) over time using exponential decay rates for mRNA (λ_RNA_) and eGFP protein (λ_Fluor_), and rates of translation (k_Trans_) and protein maturation (k_MAT_). mRNA half-lives (*t_1/2 RNA_*) were calculated from the observed mRNA decay rates. (C) Correlation between the computationally determined eGFP-degron mRNA half-lives determined by computational models that excluded (Figure 6C) or include (Figure S5B) a term for delivery of the RNA.

**Table S1:** Nearest neighbor base pairing energies for modified nucleotides, related to Figure 3. Nearest-neighbor thermodynamic parameters along with the experimentally determined error for Watson-crick base pairs containing unmodified uridine (values from (Xia et al., 1998)), Ψ, m^1^Ψ, or mo^5^U. The modified nucleotide(s) for each nearest neighbor pair is highlighted in red. Parameters were derived by linear regression of UV-melting data from X short oligonucleotides containing global substitutions, as described in (Xia et al., 1998).

| Parameter | Uridine (Xia et al., 1998) | m<sup>1</sup>Ψ | mo<sup>5</sup>U | Ψ |
| --- | --- | --- | --- | --- |
| AA/UU | -0.93 ± 0.03 | -1.18 ± 0.4 | -0.64 ± 0.07 | -1.23 ± 0.05 |
| AU/UA | -1.1 ± 0.08 | -1.13 ± 0.12 | -0.77 ± 0.09 | -1.52 ± 0.14 |
| UA/AU | -1.33 ± 0.09 | -1.86 ± 0.15 | -0.95 ± 0.11 | -1.71 ± 0.16 |
| CU/GA | -2.08 ± 0.06 | -1.83 ± 0.09 | -1.60 ± 0.07 | -2.10 ± 0.10 |
| CA/GU | -2.11 ± 0.07 | -2.26 ± 0.07 | -1.87 ± 0.06 | -2.35 ± 0.08 |
| GU/CA | -2.24 ± 0.06 | -2.43 ± 0.08 | -2.00 ± 0.06 | -2.50 ± 0.08 |
| GA/CU | -2.35 ± 0.06 | -2.67 ± 0.08 | -2.30 ± 0.07 | -2.51 ± 0.10 |

**Table S2: SHAPE-MaP reactivities, related to Figure 4.**

SHAPE-MaP reactivity data for Luc and hEpo mRNAs containing uridine, m^1^Ψ or mo^5^U indicated as (U, 1-m-pU, and 5mo-U) in Row 1. The value ‘-999’ is used to note positions of ‘NO DATA’ where low read-depth or high background mutation interfere with accurate quantification of SHAPE reactivities.

**Table S3. Table of mRNA sequences, related to Figure 1.**

A table file with the names and complete sequences for all mRNA variants of hEpo, eGFP, and Luc used in this manuscript.

## Methods

### Contact for Reagent and Resource Sharing

Further information and requests for resources and reagents should be directed to and will be fulfilled by Iain J. McFadyen

### Cell culture models

HeLa (ATCC CCL-2), Vero (ATCC CCL-81), BJ (ATCC CRL-2522), HepG2 (ATCC HB-8065), and AML12 (ATCC CRL-2254) cells were maintained in Dulbecco’s Modified Eagle’s Medium (DMEM) supplemented with GlutaMAX, HEPES, high glucose (Life Technologies, cat. no. 10564-011), 10% fetal bovine serum (FBS) (Life Technologies, cat. no. 10082-147) and sodium pyruvate (Life Technologies, cat. no. 11360-070) at 37°C in a humidified incubator at 5% CO_2_ atmosphere. Cells were passaged every 3-4 days with 0.25% trypsin-EDTA solution (Life Technologies, cat. no. 25200-056) and washed with sterile PBS (Life Technologies, cat. no. 10010-049) under aseptic conditions, for no more than 20 passages.

For all *in vitro* assays carried out in primary mouse hepatocytes, cryopreserved primary mouse hepatocytes (ThermoFisher cat. no. HMCPIS) were thawed and immediately plated for use in CHRM (ThermoFisher cat. no. CM7000), Williams Medium E supplemented with Hepatocyte Plating Supplement Pack (Serum-Containing). Plates were incubated at 37 °C in a humidified incubator at 5% CO_2_ atmosphere for 5 hours before changing media to serum free media (William’s E Maintenance Media – Without Serum). Plates were incubated at 37 °C in a humidified incubator at 5% CO_2_ atmosphere for all periods between active use.

### Mice models

*In vivo* protein expression experiments for hEpo and Luc mRNAs were performed using CD-1 and BALB/C mouse models.

### Sequence Design

eGFP variants (G_1_-G_4_) were stochastically generated using only frequently used codons. For hEpo, we obtained one mammalian codon-optimized sequence variant (E_CO_) (Welch et al., 2009) and eight variants generated by combining two unique sequences encoding the first 30 amino-acids (H_A_, H_B_) with four different variants of the remainder of the CDS (E_1_, E_2_, E_3_, E_4_). A larger set of Luc variants deterministically encoded each instance of a given amino acid throughout the coding sequence with the same single codon, with the set of 20 codons used differing between variants.

### mRNA Preparation

We selected three different proteins, human erythropoietin (hEpo), enhanced green fluorescent protein (eGFP) and firefly luciferase (Luc) and synthesized sequence variants *in vitro* using all unmodified nucleotides or **global substitutions** of uridine (U) for the modified uridine analogs pseudouridine (Ψ), N^1^-methyl-pseudouridine (m^1^Ψ), 5-methyoxy-urdine (mo^5^U), or a combination of Ψ and 5-methyl-cytidine (m^5^C) (Figure 1A). This was accomplished through *in vitro* transcription reactions in which uracil triphosphate was replaced with the triphosphate of the modified nucleotide. These proteins vary in their fundamental properties including biological function, protein structure, amino acid composition, length of coding sequence (from 579 to 1,653 nucleotides), and subcellular localization (intracellular or secreted). In all cases, the coding sequence was flanked by identical 5’ and 3’ untranslated regions (UTRs) capable of supporting high levels of protein expression (Figure 1B). Thus, total protein expression from these exogenous RNAs is determined by the combined impact of the primary coding sequence and the nucleotides used.

For simplicity and ease of analysis, we designed mRNA sequences based on simple one-to-one codon sets (i.e. each amino acid is encoded by the same codon at every instance of the amino acid, see Table S3 that disfavored the use of rare codons). Regions of increased rare codon frequency have been shown to decrease protein expression and mRNA stability (Presnyak et al., 2015; Weinberg et al., 2016). The hEpo protein contains a 9-amino acid (27 nucleotide) signal peptide sequence that is removed from the mature protein after targeting the protein to the endoplasmic reticulum (ER) for secretion. To evaluate whether codon choice had different effects in the signal peptide region, we also tested additional sequence designs for hEpo in which a leader region of 30 amino acids was encoded using two distinct codon sets: L1 (an AU-rich codon set) and L2 (a GC-rich codon set) (Figure 1C).

All mRNAs were synthesized by T7 RNA polymerase *in vitro* transcription reaction (IVT) (New England Biolabs cat. no. M0251L) and purified using standard techniques. The following combinations of nucleotides were used: all unmodified nucleotides, or unmodified adenosine, cytidine, and guanosine with pseudouridine (Ψ), N^1^-methyl-pseudouridine (m^1^Ψ), or 5-methoxy-uridine (mo^5^U), or unmodified adenosine and guanosine with pseudouridine and 5-methyl-cytidine (Ψ/m^5^C). DNA templates for IVT were generated by PCR amplification of codon-optimized sequences custom-ordered as plasmids from DNA2.0. All mRNAs were capped using the Vaccinia enzyme m^7^G capping system (New England Biolabs M2080S). All mRNA samples were analyzed for purity and cap content by capillary electrophoresis.

### mRNA Transfection

HeLa, Vero, BJ, AML12 and Primary Hepatocytes were seeded in 100uL per well of 96 well plates at a concentration of 2×10^5^ cells/mL one day prior to transfection and incubated overnight under standard cell culture conditions. For transfection, 50ng of mRNA was lipoplexed with 0.5uL Lipofectamine-2000 (ThermoFisher cat. no 11668027), brought to a volume of 20uL with Opti-MEM (ThermoFisher cat. no. 31985062) and directly added to cell media. All transfections were performed in duplicate.

### Expression Assays

Single endpoint Luc expression assays were conducted 24 hours post transfection, unless otherwise specified. The Luc Assay System (Promega cat. no. E1501) was used as per manufacturer’s suggested protocol with 100uL lysis buffer at 1:10 dilution with Luc assay reagent. Luminescence was measured on a Synergy H1 plate reader.

Single endpoint hEpo expression assays were conducted 24 hours post transfection, unless otherwise specified. The Human Erythropoietin Platinum ELISA kit (Affymetrix cat. no. BMS2035) was used as per manufacturer’s suggested protocol.

Single endpoint eGFP expression assays were conducted 24 hours post transfection, unless otherwise specified. Fluorescence was measured at an excitation wavelength of 488nm and emission wavelength of 509nm on a Synergy H1 plate reader.

Single endpoint interferon-beta (IFN-β) expression assays were conducted on cell supernatant 48 hours post transfection. The Human IFN-Beta ELISA kit (R&D Systems cat. no. 41410) was used as per manufacturer’s suggested protocol.

Luc mRNAs with m^1^Ψ and mo^5^U and a negative control mRNA lacking a poly(A) tail were electroporated into AML12 cells and both protein expression and RNA abundance was assayed at 1, 2, 3, 5, 18, and 24 hours. Half-lives were calculated using exponential decay curves. We were unable to reliably assay RNA concentration at time points earlier than 1 hour due to technical variability in the samples soon after electroporation. Luciferase expression was also determined for electroporated RNA at every hour from 1 to 6 hours post electroporation in order to ensure that delivery did not dramatically change the expression phenotype.

### In vivo studies

We measured reporter protein expression from exogenous mRNA in CD-1 and BALB/C mouse models.

Luc mRNAs were formulated in MC3 lipid nanoparticles at a concentration of 0.03mg/mL, administered intravenously to CD-1 mice at a dose of 0.15mg/kg of body mass and measured for expression by whole body Bioluminescence Imaging (BLI) at 6 hours post injection. hEpo mRNAs were formulated in MC3 lipid nanoparticles at a concentration of 0.01mg/mL, administered intravenously to BALB/C mice at a dose of (0.05mg/kg of body weight and measured for serum hEpo concentration using Human Erythropoietin Quantikine IVD ELISA kit (R&D Systems cat. no. DEP00) at 6 hours post injection.

### UV Melting

Absorbance was measured at 260nm on the Cary100 UV Vis Spectrometer as RNA, in 2mM Sodium citrate buffer (pH=6.5), was heated from 25°C to 80°C at a rate of 1°C/minute, and then cooled from 80°C to 25°C at a rate of 1°C/minute. This cycle was repeated three times in total. First derivative of absorbance values were then analyzed as a function of temperature.

### Determination of Nearest-Neighbor Thermodynamic Parameters

UV-melting experiments were performed on 39 synthetic RNA duplexes with Ψ, m^1^Ψ, and mo^5^U instead of uridine. The duplex sequences were designed to enable the full thermodynamic parameters for the nearest neighbor free energy contributions for each modified nucleotide to be determined using established methods (Xia et al., 1998).

Raw data were collected through absorbance versus duplex melting temperature profiles over six different synthetic oligonucleotide concentrations in 1M NaCl, 10mM Na_2_HPO_4_, and 0.5mM Na_2_EDTA, pH 6.98 salt buffer. These data were then processed using *Meltwin* v.3.5 to obtain a full thermodynamic parameter set through two different methods, those methods being the LnCt/4 vs. Tm^-1^ method and the Marquardt non-linear curve fit method.

### SHAPE-MaP

All purified IVT RNAs were denatured at 80°C for 3 minutes prior to analysis. After denaturation, RNAs were folded in 100mM HEPES, pH 8.0, 100mM NaCl and 10mM MgCl_2_ for 15 minutes at 37°C. All RNAs were then selectively modified with 10mM 1-methyl-6-nitroisatoic anhydride (1M6) (Sigma-Aldrich cat no. S888079-250MG) for 5 minutes at 37°C. Background (no SHAPE reagent) and denatured (SHAPE modified fully denatured RNA) controls were prepared in parallel.

After SHAPE modification, RNA was purified and fragmented using 15mM MgCl_2_ at 94°C for 4 minutes. Purified fragments were then randomly primed with N_9_ primer at 70°C for 5 minutes. Primer extension was carried out in 50mM Tris-HCl, pH 8.0, 75mM KCl, 1mM dNTPs, 5mM DTT and 6.25mM MnCl_2_ with Superscript II reverse transcriptase (ThermoFisher cat. no. 10864014) for 3 hours at 45°C. RNA-seq library prep was done with the NEBNext Ultra Directional RNA Library Prep Kit for Illumina (New England Biolabs cat. no. E7420) per the manufacturer’s standard protocol.

RNA-seq libraries were sequenced on the Illumina MiSeq using 50 cycle sequencing kit. Raw sequencing data was analyzed using the publicly available ShapeMapper software (Siegfried et al., 2014). The resulting reactivity data were analyzed using a sliding window (median SHAPE) approach to quantify the degree of structure at each position in the RNA, as has previously been described (Watts et al., 2009).

## Quantification and Statistical Analysis

### Comparison of codon effects on translation

Luc expression values from 39 Luc variants were used in 865 pairwise comparisons between synonymous codons to yield p-value testing whether inclusion of specific codons impacted protein expression by ANOVA. Graph Pad software was used to determine p-values and p-values < 0.05 were considered significant.

### Determination of structure function relationship in SHAPE data

The log normalized values for sliding window average of SHAPE reactivites from every position within the RNA were compared to the expression levels determined in HeLa cells. Linear regression was used to determine the degree of correlation between SHAPE and protein

### Computational modeling of eGFP expression data

Time-course data was collected from HeLa and AML12 cells transfected with the designed eGFP-degron mRNAs. The eGFP-degron construct reduces protein half-life to under 1 hour, so that changes in eGFP fluorescence over time directly correlate to the kinetics of translation and mRNA decay (Li et al., 1998). Fluorescence from the transfected cells was monitored over a 20-hour time course, and total active protein levels were calculated by integrating the area under the curve. These data were used to fit a the computational model of active protein production and degradation (Figure 6C) in which rate terms for protein maturation and degradation were held constant and the translation efficiency and rate of RNA degradation were allowed to vary to find the best fit to the experimental data. Data fitting was done in python using the Scikit learn module.

## Data availability

Raw sequencing files (.fastq) and processed reactivities from the SHAPE-MaP structure probing experiment are deposited into GEO under the ID codes XXXXXXXXXX.

## References

Buhr, F., Jha, S., Thommen, M., Mittelstaet, J., Kutz, F., Schwalbe, H., Rodnina, M.V., and Komar, A.A. (2016). Synonymous Codons Direct Cotranslational Folding toward Different Protein Conformations. Mol Cell 61, 341–351.

Crameri, A., Whitehorn, E.A., Tate, E., and Stemmer, W.P. (1996). Improved green fluorescent protein by molecular evolution using DNA shuffling. Nat Biotechnol 14, 315–319.

Ding, Y., Tang, Y., Kwok, C.K., Zhang, Y., Bevilacqua, P.C., and Assmann, S.M. (2014). In vivo genome-wide profiling of RNA secondary structure reveals novel regulatory features. Nature 505, 696–700.

Dominissini, D., Nachtergaele, S., Moshitch-Moshkovitz, S., Peer, E., Kol, N., Ben-Haim, M.S., Dai, Q., Di Segni, A., Salmon-Divon, M., Clark, W.C., et al. (2016). The dynamic N(1)-methyladenosine methylome in eukaryotic messenger RNA. Nature 530, 441–446.

Futcher, B., Gorbatsevych, O., Shen, S.H., Stauft, C.B., Song, Y., Wang, B., Leatherwood, J., Gardin, J., Yurovsky, A., Mueller, S., et al. (2015). Reply to Simmonds et al.: Codon pair and dinucleotide bias have not been functionally distinguished. Proc Natl Acad Sci U S A 112, E3635–3636.

Gu, W., Zhou, T., and Wilke, C.O. (2010). A universal trend of reduced mRNA stability near the translation-initiation site in prokaryotes and eukaryotes. PLoS Comput Biol 6, e1000664.

Gustafsson, C., Govindarajan, S., and Minshull, J. (2004). Codon bias and heterologous protein expression. Trends Biotechnol 22, 346–353.

Harcourt, E.M., Kietrys, A.M., and Kool, E.T. (2017). Chemical and structural effects of base modifications in messenger RNA. Nature 541, 339–346.

Hinnebusch, A.G., Ivanov, I.P., and Sonenberg, N. (2016). Translational control by 5’-untranslated regions of eukaryotic mRNAs. Science 352, 1413–1416.

Horstick, E.J., Jordan, D.C., Bergeron, S.A., Tabor, K.M., Serpe, M., Feldman, B., and Burgess, H.A. (2015). Increased functional protein expression using nucleotide sequence features enriched in highly expressed genes in zebrafish. Nucleic Acids Res 43, e48.

Jungfleisch, J., Nedialkova, D.D., Dotu, I., Sloan, K.E., Martinez-Bosch, N., Bruning, L., Raineri, E., Navarro, P., Bohnsack, M.T., Leidel, S.A., et al. (2017). A novel translational control mechanism involving RNA structures within coding sequences. Genome Res 27, 95–106.

Juszkiewicz, S., Chandrasekaran, V., Lin, Z., Kraatz, S., Ramakrishnan, V., and Hegde, R.S. (2018). ZNF598 Is a Quality Control Sensor of Collided Ribosomes. Mol Cell 72, 469–481 e467.

Kariko, K., Buckstein, M., Ni, H., and Weissman, D. (2005). Suppression of RNA recognition by Toll-like receptors: the impact of nucleoside modification and the evolutionary origin of RNA. Immunity 23, 165–175.

Kariko, K., Muramatsu, H., Welsh, F.A., Ludwig, J., Kato, H., Akira, S., and Weissman, D. (2008). Incorporation of pseudouridine into mRNA yields superior nonimmunogenic vector with increased translational capacity and biological stability. Mol Ther 16, 1833–1840.

Kertesz, M., Wan, Y., Mazor, E., Rinn, J.L., Nutter, R.C., Chang, H.Y., and Segal, E. (2010). Genome-wide measurement of RNA secondary structure in yeast. Nature 467, 103–107.

Kierzek, E., and Kierzek, R. (2003). The thermodynamic stability of RNA duplexes and hairpins containing N6-alkyladenosines and 2-methylthio-N6-alkyladenosines. Nucleic Acids Res 31, 4472–4480.

Kimchi-Sarfaty, C., Oh, J.M., Kim, I.W., Sauna, Z.E., Calcagno, A.M., Ambudkar, S.V., and Gottesman, M.M. (2007). A “silent” polymorphism in the MDR1 gene changes substrate specificity. Science 315, 525–528.

Kormann, M.S., Hasenpusch, G., Aneja, M.K., Nica, G., Flemmer, A.W., Herber-Jonat, S., Huppmann, M., Mays, L.E., Illenyi, M., Schams, A., et al. (2011). Expression of therapeutic proteins after delivery of chemically modified mRNA in mice. Nat Biotechnol 29, 154–157.

Li, X., Zhao, X., Fang, Y., Jiang, X., Duong, T., Fan, C., Huang, C.C., and Kain, S.R. (1998). Generation of destabilized green fluorescent protein as a transcription reporter. J Biol Chem 273, 34970–34975.

Lu, Z.J., Turner, D.H., and Mathews, D.H. (2006). A set of nearest neighbor parameters for predicting the enthalpy change of RNA secondary structure formation. Nucleic Acids Res 34, 4912–4924.

Mortimer, S.A., Kidwell, M.A., and Doudna, J.A. (2014). Insights into RNA structure and function from genome-wide studies. Nat Rev Genet 15, 469–479.

Mustoe, A.M., Busan, S., Rice, G.M., Hajdin, C.E., Peterson, B.K., Ruda, V.M., Kubica, N., Nutiu, R., Baryza, J.L., and Weeks, K.M. (2018). Pervasive Regulatory Functions of mRNA Structure Revealed by High-Resolution SHAPE Probing. Cell 173, 181–195 e118.

Nakamura, Y., Gojobori, T., and Ikemura, T. (2000). Codon usage tabulated from international DNA sequence databases: status for the year 2000. Nucleic Acids Res 28, 292.

Newby, M.I., and Greenbaum, N.L. (2001). A conserved pseudouridine modification in eukaryotic U2 snRNA induces a change in branch-site architecture. RNA 7, 833–845.

Plotkin, J.B., and Kudla, G. (2011). Synonymous but not the same: the causes and consequences of codon bias. Nat Rev Genet 12, 32–42.

Pop, C., Rouskin, S., Ingolia, N.T., Han, L., Phizicky, E.M., Weissman, J.S., and Koller, D. (2014). Causal signals between codon bias, mRNA structure, and the efficiency of translation and elongation. Mol Syst Biol 10, 770.

Presnyak, V., Alhusaini, N., Chen, Y.H., Martin, S., Morris, N., Kline, N., Olson, S., Weinberg, D., Baker, K.E., Graveley, B.R., et al. (2015). Codon optimality is a major determinant of mRNA stability. Cell 160, 1111–1124.

Ramani, V., Qiu, R., and Shendure, J. (2015). High-throughput determination of RNA structure by proximity ligation. Nat Biotechnol 33, 980–984.

Rauscher, R., and Ignatova, Z. (2018). Timing during translation matters: synonymous mutations in human pathologies influence protein folding and function. Biochem Soc Trans 46, 937–944.

Sabnis, S., Kumarasinghe, E.S., Salerno, T., Mihai, C., Ketova, T., Senn, J.J., Lynn, A., Bulychev, A., McFadyen, I., Chan, J., et al. (2018). A Novel Amino Lipid Series for mRNA Delivery: Improved Endosomal Escape and Sustained Pharmacology and Safety in Non-human Primates. Mol Ther.

Scherer, S., and Cold Spring Harbor Laboratory. Press. (2010). Guide to the human genome (Cold Spring Harbor, N.Y.: Cold Spring Harbor Laboratory Press).

Schoenberg, D.R. (2011). Mechanisms of endonuclease-mediated mRNA decay. Wiley Interdiscip Rev RNA 2, 582–600.

Shah, P., Ding, Y., Niemczyk, M., Kudla, G., and Plotkin, J.B. (2013). Rate-limiting steps in yeast protein translation. Cell 153, 1589–1601.

Siegfried, N.A., Busan, S., Rice, G.M., Nelson, J.A., and Weeks, K.M. (2014). RNA motif discovery by SHAPE and mutational profiling (SHAPE-MaP). Nat Methods 11, 959–965.

Simmonds, P., Tulloch, F., Evans, D.J., and Ryan, M.D. (2015). Attenuation of dengue (and other RNA viruses) with codon pair recoding can be explained by increased CpG/UpA dinucleotide frequencies. Proc Natl Acad Sci U S A 112, E3633–3634.

Simms, C.L., Yan, L.L., and Zaher, H.S. (2017). Ribosome Collision Is Critical for Quality Control during No-Go Decay. Mol Cell 68, 361–373 e365.

Spitale, R.C., Flynn, R.A., Zhang, Q.C., Crisalli, P., Lee, B., Jung, J.W., Kuchelmeister, H.Y., Batista, P.J., Torre, E.A., Kool, E.T., et al. (2015). Structural imprints in vivo decode RNA regulatory mechanisms. Nature 519, 486–490.

Tuller, T., Carmi, A., Vestsigian, K., Navon, S., Dorfan, Y., Zaborske, J., Pan, T., Dahan, O., Furman, I., and Pilpel, Y. (2010). An evolutionarily conserved mechanism for controlling the efficiency of protein translation. Cell 141, 344–354.

Tuller, T., and Zur, H. (2015). Multiple roles of the coding sequence 5' end in gene expression regulation. Nucleic Acids Res 43, 13–28.

Tulloch, F., Atkinson, N.J., Evans, D.J., Ryan, M.D., and Simmonds, P. (2014). RNA virus attenuation by codon pair deoptimisation is an artefact of increases in CpG/UpA dinucleotide frequencies. Elife 3, e04531.

Wan, Y., Qu, K., Zhang, Q.C., Flynn, R.A., Manor, O., Ouyang, Z., Zhang, J., Spitale, R.C., Snyder, M.P., Segal, E., et al. (2014). Landscape and variation of RNA secondary structure across the human transcriptome. Nature 505, 706–709.

Watts, J.M., Dang, K.K., Gorelick, R.J., Leonard, C.W., Bess, J.W., Jr., Swanstrom, R., Burch, C.L., and Weeks, K.M. (2009). Architecture and secondary structure of an entire HIV-1 RNA genome. Nature 460, 711–716.

Weinberg, D.E., Shah, P., Eichhorn, S.W., Hussmann, J.A., Plotkin, J.B., and Bartel, D.P. (2016). Improved Ribosome-Footprint and mRNA Measurements Provide Insights into Dynamics and Regulation of Yeast Translation. Cell Rep 14, 1787–1799.

Welch, M., Govindarajan, S., Ness, J.E., Villalobos, A., Gurney, A., Minshull, J., and Gustafsson, C. (2009). Design parameters to control synthetic gene expression in Escherichia coli. PLoS One 4, e7002.

Wen, J.D., Lancaster, L., Hodges, C., Zeri, A.C., Yoshimura, S.H., Noller, H.F., Bustamante, C., and Tinoco, I. (2008). Following translation by single ribosomes one codon at a time. Nature 452, 598–603.

Wilamowski, M., Gorecki, A., Dziedzicka-Wasylewska, M., and Jura, J. (2018). Substrate specificity of human MCPIP1 endoribonuclease. Sci Rep 8, 7381.

Xia, T., SantaLucia, J., Jr., Burkard, M.E., Kierzek, R., Schroeder, S.J., Jiao, X., Cox, C., and Turner, D.H. (1998). Thermodynamic parameters for an expanded nearest-neighbor model for formation of RNA duplexes with Watson-Crick base pairs. Biochemistry 37, 14719–14735.

Zarai, Y., Margaliot, M., and Tuller, T. (2016). On the Ribosomal Density that Maximizes Protein Translation Rate. PLoS One 11, e0166481.

